# *ZGRF1* variants implicated in a sensory reflex epilepsy

**DOI:** 10.1101/728188

**Authors:** Shalini Roy Choudhury, Parthasarathy Satishchandra, Sanjib Sinha, Anuranjan Anand

## Abstract

Hot water epilepsy (HWE) is a sensory reflex epilepsy in which seizures are precipitated by a stimulus such as contact with hot water. While there is a genetic basis to its etiology, identity of the genes underlying this relatively uncommon disorder has remained unknown. Here, we present the results of our studies aimed at identifying a causative gene in a south Indian four-generation family with several affected members. We conducted whole-exome sequencing and examined a known locus that maps to 4q24-q28 (*HWE2*, MIM: 613340) that we had previously identified. We identified a sequence variant, c.1805C>T (p.Thr602Ile) in *ZGRF1*, located within the *HWE2* locus, co-segregating with the disorder. The transcript structure of *ZGRF1* was examined in 288 HWE patients, and five additional missense variants, Arg326Gln, Glu660Gly, Arg1862*, Phe1940Leu and Asp1984Gly, present exclusively or almost exclusively in the patients were found. Functional correlates of the six variants identified were examined in cultured mammalian cells. We observed spindle pole defects during cell division and partially disrupted localization of UPF1, a protein involved in cell cycle regulation, at the spindles. Our observations provide insights into the genetic basis of HWE and suggest the involvement of hitherto unanticipated molecular process in this disorder.

## Introduction

Sensory reflex epilepsy comprises a class of rare human epilepsies (Italiano et al., 2016) in which seizures are triggered by a wide range of sensory stimuli. A somatosensory stimulus of contact with hot water at temperatures in the range 40-50°C is known to trigger onset of seizures in a condition known as hot water epilepsy (HWE). This disorder has also been described as water immersion epilepsy or bathing epilepsy. HWE was first reported from New Zealand (Allen, 1945), following which there have been reports from different parts of the world: Australia (Keipert, 1969), the United States (Stensman and Ursing, 1971), India (Mani et al., 1974; Satishchandra et al., 1988; Satishchandra, 2003), the United Kingdom (Moran, 1976), Canada (Szymonowicz and Meloff, 1978), Japan (Kurata, 1979; Morimoto et al., 1985) and Turkey (Bebek et al., 2001). Most of the HWE cases so far reported are from India. HWE is prevalent in the southern parts of the country where hot water bathing after applying hot oil on the head is a common cultural custom. The seizures are often accompanied by a dazed look, state of confusion, indistinct speech, epigastric sensations and compulsive behavior (Satishchandra, 2003). Individuals with HWE exhibit psychiatric symptoms during the phase of aura, with a sense of fear, visual and auditory hallucinations and déjà vu, and experience feelings of living in the future. About 10% of the patients are habituated to self-induction of seizures with a sense of intense desire and pleasure, provoking these patients to continuously pour hot water over their head to induce seizures (Satishchandra et al., 1988). Both temperature of water and contact sensation seem to be important for the phenotypic manifestation of this disorder.

While the first case of HWE was reported as early as 1945 (Allen, 1945), a genetic basis for this disorder is only now beginning to be examined. Familial clustering and positive family histories have been reported for 7-15% of the patients from India (Mani et al., 1974, Satishchandra et al., 1988) and for 10% of the patients from Turkey (Bebek et al., 2001). Three HWE loci are known which map to 10q21.3-q22.3 (*HWE1*, MIM: 613339; Ratnapriya et al., 2009a), 4q24-q28 (*HWE2*, MIM: 613340; Ratnapriya et al., 2009b) and 9p24.3-p23 (Karan et al., 2018). Mutations in *SYN1* have been reported in a French-Canadian family with X-linked focal epilepsy and reflex bathing seizures (Nguyen et al., 2015). A loss-of-function mutation in *GPR56* has been identified in a 5-year-old Portuguese boy with bilateral frontoparietal polymicrogyria who presented with HWE (Santos-Silva et al., 2015). Partial loss-of-function at *SLC1A1* has been reported in HWE families from India (Karan et al., 2017). We examined the 4q24-q28 (*HWE2*) locus to identify a candidate gene for epilepsy in a four-generation family from south India. Here, we report our studies that indicate the involvement of *ZGRF1* (zinc finger GRF-type containing 1) variants in HWE.

## Subjects and methods

### Family ascertainment and clinical characterization

Hot water epilepsy families were ascertained at the Department of Neurology, National Institute of Mental Health and Neurosciences (NIMHANS), Bangalore. Family 227 is a four-generation family of 28 members, of which 10 were affected, three clinically asymptomatic and 15 unaffected (Figure 1A). HWE in the family is transmitted in an apparently autosomal dominant mode with incomplete penetrance. Clinical diagnoses were made by a qualified neurologist according to ILAE guidelines for epilepsy classification (Engel J, 2001) and clinical criterion for seizures upon hot water bath based on clinical histories. Information regarding seizures and family histories of epilepsy or other neurological disorders was obtained from patients and family members who had witnessed such episodes in the affected individuals. Family 227 was recruited through proband III:7, diagnosed with HWE at 20 years of age. His seizure episodes were characteristic of HWE and his inter-ictal EEG showed discharges arising in the left anterior temporal region. No hippocampal abnormality was detected. He was treated with intermittent prophylactic clobazam therapy. Two individuals, II:7 and IV:1, had histories of febrile convulsions (FCs). Individuals II:2, II:3 and III:2 were asymptomatic but had at least one child manifesting HWE. Peripheral blood samples were collected from each participating individual and DNA was isolated from white blood cells using a standard phenol-chloroform method. This study had the approval of the Institutional Human Bioethics Committees of NIMHANS and JNCASR. All members of the family who participated in the study provided informed written consent.

**Figure 1:**
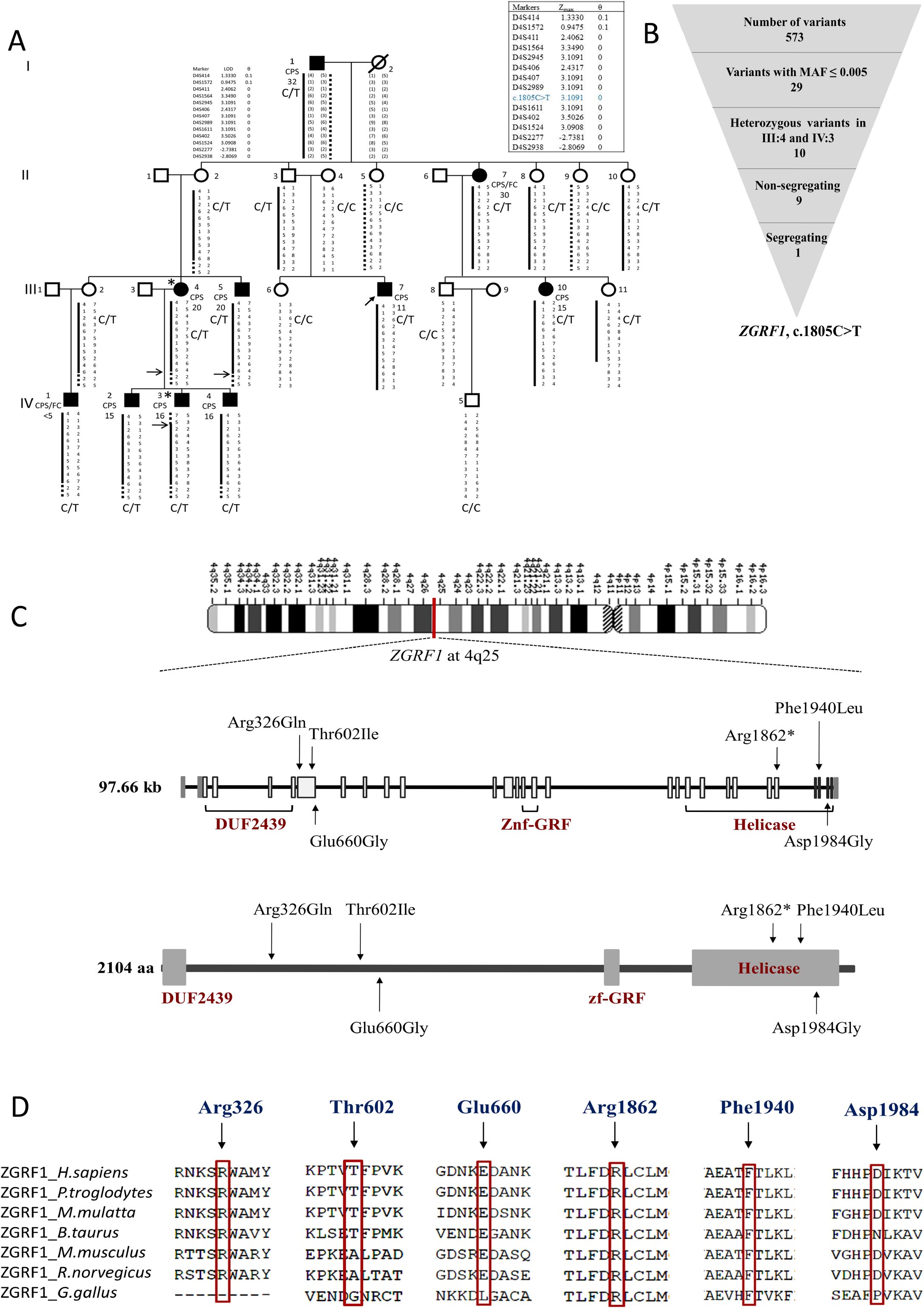
*ZGRF1* at 4q24-q28. (A) Pedigree of Family 227 showing the segregation of the disease haplotype and c.1805C>T. Individuals (lll:4 and lV:3) marked with an asterisk(*) were whole-exome sequenced (modified from Ratnapriya et al., 2009b). The maximum LOD scores obtained for the markers and for the variant are shown in the box. (B) Workflow for screening and filtering the gene variants in the 4q24-q28 target region. (C) Chromosomal location, gene structure, and protein structure of ZGRF1 showing the positions of the six rare variants. (D) Amino acid conservation of the six residues across certain species.

### DNA sequencing and analysis

Whole exome sequencing was carried out for III:4 and IV:3, an affected parent-offspring pair using the Agilent SureSelect Human All Exon 50Mb Kit (Agilent Technologies, Santa Clara, USA). Genomic DNA sequencing libraries were prepared according to manufacturer’s protocol (Agilent). Briefly, genomic DNA was fragmented, purified, analysed for size distribution and concentration, adapters were ligated, hybridized with probes and enriched for bound regions. The libraries were sequenced for 100-bp paired-end reads on Illumina Genome Analyzer (GAIIx). Reads with at least 70% of the bases having a minimum Phred score of 20 were obtained by SeqQC-V2.0. FASTQ files were trimmed and aligned to the reference human genome (hg19, GRCh37) using Burrows-Wheeler Aligner (BWA) 0.6.0 (Li and Durbin, 2009). Duplicate reads and those with inappropriate read pair were removed using SAMtools v0.1.7a (Li et al., 2009). Alignment statistics were obtained from BEDTools v-2.12.0 (Quinlan and Hall, 2010). Variants were called by SAMtools at a Phred-like SNP quality score of ≥20 and annotated against dbSNP131 and dbSNP135. Sequence reads at the locus were manually examined and updated control frequencies of annotated variants were confirmed in dbSNP144, 1000 Genomes, Exome Variant Server (ESP), and The Genome Aggregation Database (gnomAD). About 8% of the 4q24-q28 locus not covered by exome sequencing was Sanger sequenced using specific primer pairs. Common variants with a minor allele frequency (MAF>0.005) across databases within the target region 4q24-q28 (between D4S1572 and D4S2277) were excluded from further analysis. Variants with MAF≤0.005 present in the exons, flanking introns (±100bp) and UTRs were examined further. Rare, heterozygous variants were confirmed by Sanger sequencing, examined for segregation in the family, and coded according to HGVS (Human Genome Variation Society).

### Mutation analysis of *ZGRF1* in HWE patients

The complete transcript of *ZGRF1* (RefSeq NM_018392.4) comprising 27 coding exons, flanking introns, 5’UTR and 3’UTR was amplified in 288 HWE patients and ethnicity-matched controls using an ABI9700 Thermal Cycler (Applied Biosystems, Foster City, USA). The amplified products were sequenced using BigDye™ Terminator Cycle Sequencing v3.0 reagents (Applied Biosystems) and were analysed on an ABI3730 Genetic Analyzer (Applied Biosystems).

### Mutagenesis

The full-length cDNA of *ZGRF1* cloned into mammalian pCMV6 vector with C-terminal Myc-DDK tag (RC229463, Origene) was used as the template for introducing six *ZGRF1* variants: Arg326Gln, Thr602Ile, Glu660Gly, Arg1862*, Phe1940Leu and Asp1984Gly. Mutagenesis was performed using the QuikChange II XL Site-Directed Mutagenesis kit (Agilent). Plasmids with the *ZGRF1* variants were purified using QIAprep Spin Miniprep columns (Qiagen, Hilden, Germany) and confirmed by Sanger sequencing (For a complete list of the primers used for *ZGRF1* sequencing, see Supporting Tables S1, S2 and S3).

### Reverse-transcription polymerase chain reaction

Brain cDNA from cerebral cortex, cerebellum, hippocampus and hypothalamus (Takara Bio, Santa Clara, USA) was used to examine the expression of *ZGRF1* transcriptsin the human brain. Two primer pairs5’-TCCAAAGACACAGAAGCACA-3’ and 5’-ACCTGC AAGAAGTCAATCTGC-3’, and 5’-TCAGCCTAGGAGCAACATTGA-3’ and 5’-TCCAA CTACTCGAACCTGCT-3’ were used to amplify cDNA from the N-terminal and C-terminal regions of *ZGRF1*, respectively. The amplified products were purified using Qiaquick columns (Qiagen) and sequence confirmed.

### Cell culture and transfection

HEK293 (ATCC, Manassas, USA), SH-SY5Y (ATCC) and C6 glioma (ECACC, Sigma-Aldrich, St. Louis, USA) cells were cultured in DMEM supplemented with 10% heat-inactivated fetal bovine serum for up to 10 passages. For protein overexpression studies in HEK293 cells, transfection was carried out with 4.5 µg of plasmid using the calcium phosphate method. Cells were seeded in 35-mm dishes (for protein immunoblot analysis) or poly-L-lysine-coated coverslips (for immunocytochemistry assays). Cells were seeded 24 h before transfection, and experiments were performed 48 h after transfection.

### Immunocytochemistry

Cells were washed with phosphate-buffered saline (PBS), fixed with 2% paraformaldehyde or methanol for 15 min, permeabilized with 0.1% TritonX-100 for 10 min, incubated with blocking solution (5% BSA in PBS) for 4 h at 4°C, and stained overnight at 4°C with primary antibodies. Primary antibodies used in these experiments were: (1) rabbit polyclonal anti-ZGRF1 (1:100 for endogenous expressions, 1:1000 for overexpression studies; ab122126, Abcam, UK), (2) mouse monoclonal anti-γ-tubulin (1:5000; T5326, Sigma-Aldrich), and (3) rabbit monoclonal anti-UPF1 Alexa Fluor-488 (1:100; ab201761, Abcam). The cells were washed with PBS and incubated with secondary antibodies conjugated with Alexa Fluor-568 or Alexa Fluor-488 (Invitrogen Molecular probes, Waltham, USA) for 4 h at 4°C. Cells were then washed with PBS and counterstained with DAPI (Sigma-Aldrich). For cell-cycle synchronization experiments, HEK293 cells were incubated in culture medium containing 2 mM thymidine (Sigma-Aldrich) for 14 h, followed by mitotic enrichment in medium containing 24 μM deoxycytidine (Sigma-Aldrich) for 9 h. Three hundred transfected, mitotic cells were counted for the wild-type and six variant alleles of *ZGRF1*. Coverslips were mounted in 70% glycerol (Merck Millipore, Burlington, USA) and imaged on an LSM 880 Meta confocal laser scanning microscope (Carl Zeiss, Oberkochen, Germany) under a 63x/1.4 oil immersion objective. Images were processed and scale bars were added in Zen (Carl Zeiss). All statistical comparisons were done using GraphPad Prism 5. One-way ANOVA followed by Dunnett’s test for multiple comparisons were used for statistical analysis between the wild-type and variants. Results are shown as mean ± standard error of the mean (SEM). Differences between groups were considered statistically significant for *P* ≤ 0.05 and *P* ≤ 0.001.

### Immunoblot analysis

Cells were lysed in 200 μl of RIPA lysis buffer (50 mM Tris-HCl pH 7.4, 150 mM NaCl, 1 mM EDTA, 0.5% sodium deoxycholate, 1% NP-40 and 0.1% SDS) supplemented with protease inhibitor cocktail (Sigma-Aldrich) for 3 h at 4°C in a rotating mixer. Lysed cells were spun down at 14,000 g for 20 min at 4°C and protein concentration was determined using the bicinchoninic acid assay (Sigma-Aldrich). Approximately 50 μg of protein was loaded on an 8% polyacrylamide gel along with pre-stained protein ladder (Bio-Rad Laboratories, Hercules, USA). Proteins from the gel were electro-transferred to 0.45 μm PVDF membrane (Merck Millipore) using a wet electroblotting system (Bio-Rad Laboratories) at a constant voltage of 100 V for 1 h in the transfer buffer (25 mM Tris-HCl, 192 mM glycine, 20% methanol and 0.036% SDS) at 4°C. The membrane was blocked with 5% skimmed milk (HiMedia, Mumbai, India) for 4 h at 4°C, incubated overnight at 4°C with primary antibodies rabbit polyclonal anti-ZGRF1 (1:300; ab122126) or anti-γ-tubulin (1:5000; T5326). The blot was washed in PBS, incubated for 1 h at room temperature with rabbit or mouse horseradish peroxidase (HRP)-conjugated secondary antibodies (1:5000; Bangalore GeNei, Bangalore, India) and washed with PBS containing 0.1% Tween-20. Protein bands were detected using an enhanced chemiluminescent substrate for HRP (West Pico, Pierce, Waltham, USA). Lysates (50 μg) from different regions of the brain, amygdala, basal ganglia, cerebellum, hippocampus and hypothalamus were loaded on a 5-15% gradient gel, transferred on to a 0.45 μm nitrocellulose membrane (GE Healthcare Life Sciences, Marlborough, USA) using a semi-dry transfer system (Bio-Rad Laboratories), and the membrane was processed as above.

### Bioinformatic analysis

The complete nucleotide sequence of *ZGRF1* is available at NCBI (Accession Number NC_000004.11, assembly GRCh37, annotation release 105). It comprises 97.66 kb of sequence length spanning bases 113,460,489-113,558,151 (GRCh37 co-ordinates). Primers used in this study were designed using Primer3. Nucleotide and protein sequences across different species were obtained from NCBI HomoloGene database. Sequence conservation and multiple sequence alignment analyses were carried out using Clustal Omega. Potential pathogenic effects of the variants were predicted using SIFT (Kumar et al., 2009), PolyPhen-2 (Adzhubei et al., 2010), MutationTaster (Schwarz et al., 2010), and FATHMM (Shihab et al., 2013). pBLAST (protein-protein BLAST) was used to determine ZGRF1 homologs and DELTA-BLAST (Domain Enhanced Lookup Time Accelerated BLAST; Boratyn et al., 2012) was used to predict the domain architecture of ZGRF1. Domain-wise identities and similarities were determined by pairwise alignment using EMBOSS Needle.

## Results

### Rare variant prioritization

We performed whole exome sequencing on two affected individuals, III:4 and IV:3, from the family. For both individuals, we obtained a mapping efficiency of 95.7%, and 92% coverage of the exonic regions with ≥10 reads, at the 4q24-q28 locus (Supporting Tables S4 and S5). Employing a combination of exome-based and Sanger-based sequencing, we identified 573 variants at the target region, in the two individuals (Figure 1B). Among these, 544 were relatively common variants (MAF>0.005) across the different control databases. For remaining 29 rare variants (MAF ≤ 0.005, Supporting Tables S6 and S7) we confirmed their presence/absence and zygosity in the two individuals whose exomes were sequenced. Nineteen of the 29 variants were either homozygous, or heterozygous and present in only one of the affected members. These variants were not considered further. The remaining 10 variants were heterozygous and present in both III:4 and IV:3, and were examined for segregation in the family. Nine of these 10 variants did not segregate. We found one non-synonymous, heterozygous variant, c.1805C>T located in exon 5 of the *ZGRF1* (zinc finger GRF-type containing 1) gene, to co-segregate with the disorder in the family. *ZGRF1* c.1805C>T leads to amino acid change p.Thr602Ile in the ZGRF1 protein (Figure 1C). Among the available ZGRF1 protein sequences, this residue is conserved among higher mammals, namely *Homo sapiens, Macaca mulatta, Bos taurus* and *Pan troglodytes* (Figure 1D).

*ZGRF1* encodes a transcript of 6652 bases and a protein of 2104 amino acids. According to HGNC (HUGO Gene Nomenclature Committee), ZGRF1 belongs to the family of zinc-finger GRF-type and UPF1-like RNA helicases. There are at least six additional GRF-type proteins, and 11 proteins that belong to the UPF1-like RNA helicases family. These are spliceosomal factors, DNA/RNA helicases or nucleases. *In-silico* analyses of the ZGRF1 sequence identified three main domains in the protein (Supporting Figure S8a). An N-terminal *first domain* DUF2439 (domain of unknown function 2439) encompasses 70 amino acids. Orthologs of this domain are found in *Schizosaccharomyces pombe, Saccharomyces cerevisiae* and *Arabidopsis thaliana* (Polakova et al., 2016). This domain has been implicated in meiotic chromosome segregation, crossover recombination and telomere maintenance (Polakova et al., 2016; Silva et al., 2016). A centrally placed *second domain zf*-GRF (zinc finger-GRF type) is a presumed zinc-binding domain of 45 amino acids named after three conserved residues within the domain. This domain is present in proteins with helicase, nuclease, polymerase or topoisomerase activities. The C-terminus has the *third domain* comprising four approximately overlapping domains with predicted helicase activities, which we collectively refer to as the helicase domain. These are the AAA_11, AAA_12/UvrDC2, DNA2 and recD domains. AAA_11 and AAA_12/UvrDC2 span 231 and 185 amino acids, respectively. They belong to a large and functionally diverse AAA superfamily (ATPase Associated with diverse cellular Activities) of ring-shaped P-loop NTPases that function as molecular chaperones (Iyer et al., 2004) through energy-dependent unfolding of macromolecules (Frickey and Lupas, 2004). The DNA2 domain of 319 amino acids belongs to the superfamily I (SF1) DNA and/or RNA helicases. Structural data are available for a single SF1 RNA helicase, UPF1 (up-frameshift 1), a protein involved in nonsense-mediated mRNA decay (Cheng et al., 2007). Biochemical analysis of SF1 RNA helicases has revealed dual RNA-stimulated and DNA-stimulated ATPase and helicase activities (Raney et al., 2013). The recD domain spanning 44 amino acids is present in DNA helicases.

### *ZGRF1* variants in additional HWE patients

We examined the complete transcript structure of *ZGRF1* among 288 HWE patients. In this analysis (Table 1), we identified six rare missense variants in the gene: c.977G>A (p.Arg326Gln), c.1805C>T (p.Thr602Ile) and c.1979A>G (p.Glu660Gly) located in the long non-domain region towards the N-terminus of the protein placed between the DUF2439 and zf-GRF domains; and three variants c.5584C>T (p.Arg1862*), c.5818T>C (p.Phe1940Leu) and c.5951A>G (p.Asp1984Gly) within the helicase domain, towards the C-terminus (Figure 1C, Supporting Figure S8b). These six non-synonymous variants were either absent or rare among 576 population-matched control chromosomes of individuals from the southern parts of India, as well as among the four variant databases dbSNP, 1000 Genomes, ESP and gnomAD. The amino acid residues Arg1862 and Phe1940 are well conserved across different species examined, Glu660 and Asp1984 are conserved among all species except *G*.*gallus* and Arg326 is conserved among all species and absent in *Gallus gallus* (Figure 1D). We also found 29 single-nucleotide polymorphisms in the *ZGRF1* gene that were common among population-matched controls and in databases (Supporting Table S9).

**Table 1:** *ZGRF1* variants among HWE patients.

| Nucleotide change | Amino acid change | rsID | Occurrence in patients (n=288) | MAF in control individuals (n=288) | MAF in dbSNP 144,1000G, ESP or ExAC | Conservation score<sup>a</sup> | SIFT score<sup>b</sup> | PolyPhen-2 Hum Varscore<sup>c</sup> | Mutation-Taster probability value<sup>d</sup> | FATHMM score<sup>e</sup> |
| --- | --- | --- | --- | --- | --- | --- | --- | --- | --- | --- |
| c.977G>A | p.Arg326Gln | rs535674967 | 1 | 0 | 0.0002 (dbSNP), 0.0002 (1000G), 0.00004 (gnomAD) | 0.015 | Tolerated 0.26 | Probably damaging 0.998 | Polymorphism 0.923 | Damaging -1.96 |
| c.1805C>T | p.Thr602Ile | rs201904239 | 3 | 0.0034 | 0.0012 (dbSNP), 0.001 (1000G), 0.0008 (gnomAD) | 0.601 | Damaging 0.03 | Benign 0.012 | Polymorphism 0.999 | Damaging -1.64 |
| c.1979A>G | p.Glu660Gly | - | 1 | 0 | - | -0.817 | Damaging 0.05 | Benign 0.013 | Polymorphism 0.999 | Damaging -2.02 |
| c.5584C>T | p.Arg1862* | rs767296222 | 2 | 0 | 0.00004 (gnomAD) | -1.202 | Damaging 0.01 | - | Disease causing 0.999 | - |
| c.5818T>C | p.Phe1940Leu | rs773566516 | 2 | 0 | 0.00001 (gnomAD) | -1.094 | Damaging0 | Benign 0.284 | Disease causing 0.967 | Damaging -2.86 |
| c.5951A>G | p.Asp1984Gly | rs780748235 | 1 | 0.0017 | 0.00004 (gnomAD) | 1.562 | Tolerated 0.35 | Benign 0.108 | Polymorphism 0.999 | Tolerated -1.13 |

(RefSeq id: NM\_018392.4, CCDS3700.2)
<sup>a</sup>TheConsurf Server; conservation scores for the amino acids based on the phylogenetic relationship from 41 homologous sequences (Ashkenazy et al., 2010).
<sup>b</sup>Sorting Intolerant From Tolerant [SIFT] (Kumar et al., 2009): SIFT predicts effects for amino acid substitutions with scores ranging from 0 to 1. Variant scores ranging from 0.0 to 0.05 are considered deleterious, with scores closer to 0.0 being confidently predictive of deleteriousness. Scores between 0.05 to 1.0 are considered tolerated, and scores closer to 1.0 suggest that the variant is tolerable.
<sup>c</sup>PolymorphismPhenotyping v2 [PolyPhen-2] (Adzhubei et al., 2010): PolyPhen predicts effects for amino acid substitutions with scores ranging from 0 to 1 as being benign to damaging.
<sup>d</sup>MutationTaster (Schwarz et al 2010): Probability of prediction; values close to 1 indicate high security of the prediction.
<sup>e</sup>Functional Analysis through Hidden Markov Models (FATHMM) (Shihab et al., 2013): Combines sequence conservation with hidden Markov models to determine 'pathogenicity weights'.

### *ZGRF1* expression in the human brain

*ZGRF1* is expressed in human brain regions. We amplified ZGRF1 transcripts from Marathon-Ready™ full-length brain cDNAs from human cerebral cortex, cerebellum, hippocampus and hypothalamus. The gel bands (Figure 2A) corresponded to the expected sizes of 656 bases and 572 bases for the N-terminal ad C-terminal regions, respectively. We extracted the bands and confirmed them by sequencing. Western blot analysis indicated immunoreactivity to the anti-ZGRF1 antibody in amygdala, basal ganglia, cerebellum, hippocampus and hypothalamus of the human brain (Figure 2B). The *ZGRF1* transcripts are reported to be ubiquitously present across different human tissues (Human Protein Atlas RNA-sequencing data).

**Figure 2:**
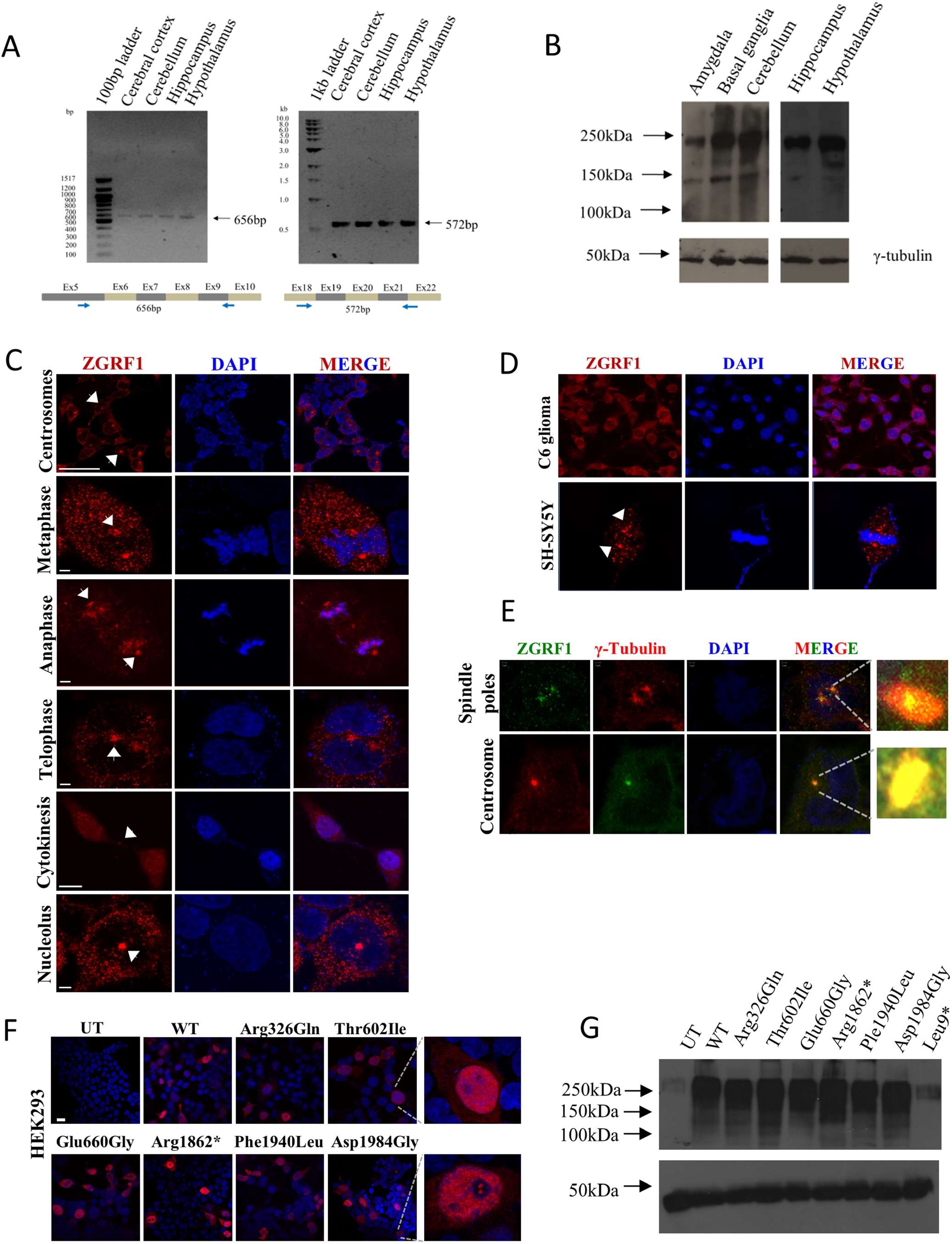
Expression of ZGRF1. (A) Expression of *ZGRF1* transcripts in different regions of the human brain. (B) ZGRF I protein expression in different human brain regions. (C) Endogenous localization of ZGRFI in cultured HEK293 cells. Arrows indicate specific localizations of the protein. Scale bars = 20 µm, 5 µm. (D) Endogenous locali zation of ZGRF I in C6 glioma and SH-SYSY cells. Arrowheads point towards spindle poles. (E) Co-localization of ZGRF I with γ-tubulin at spindle poles and centrosomes. (F) Localization of overexpressed ZGRF1 wild-type and mutant proteins in the nucleus of HEK293. Scale bar= 20 µm. (G) Wildtype (WT) and variant proteins in untransfected (UT) and overexpressing HEK293 cells.

### ZGRF1 expression during mitotic stages

In cultured HEK293 cells, ZGRF1 showed localization in the cell cytoplasm. We observed that the protein has a distinctive localization pattern in mitotic cells. ZGRF1 localizes at the centrosomes during prophase and at the spindle poles during metaphase and anaphase stages (Figure 2C). The protein localized to midbodies during telophase and cytokinesis stages. Some cells also showed the protein in the nucleolus. In SH-SY5Y neuroblastoma and rat C6 glioma cells, the protein localized in a similar manner in the cytoplasm and could be detected at the spindles poles in dividing cells (Figure 2D). ZGRF1 co-localized with γ-tubulin, a microtubule and centrosomal marker, at the spindle poles and centrosomes (Figure 2E). In immunocytochemistry experiments aimed at determining the cellular localization pattern of the ZGRF1 variant proteins (Figure 2F), wild-type ZGRF1 localized in a heterogeneous manner, majorly in the nucleus, occasionally in the cytoplasm, or both in the nucleus and cytoplasm, in HEK293 cells transiently overexpressing the gene. The ZGRF1 variant proteins localized in a similar manner as the wild-type, with no apparent disturbances in the localization patterns, or in cellular morphological features. We estimated the steady-state protein expression levels in whole-cell lysates for the wild-type and variant proteins (Figure 2G). While ZGRF1 could be detected at the endogenous level in untransfected HEK293 cells, the steady state protein level was high in cells transfected with the *ZGRF1* wild-type allele. The six ZGRF1 variant proteins showed no difference in the expression levels compared to the wild-type protein.

### *ZGRF1* variants and mitotic defects

We observed that the synchronized cultured cells with ZGRF1 variant proteins exhibited significant mitotic defects compared to those with the ZGRF1 wild-type. These defects include multipolar cells with increased number of centrosomes, disoriented spindle poles and abnormal cytokinesis (Figure 3A). The highest extent of defects (Figure 3B) was observed for Thr602Ile (73.96%) and Asp1984Gly showed 70.8% defective cells and Phe1940Leu, 69.32%. Comparable levels of defects were observed for the other variants, Glu660Gly (64.66%), Arg1862* (61%) and Arg326Gln (60.33%). We observed mitotic defects in cells overexpressing the wild-type protein (30.66%) compared to untransfected cells with endogenous expression. This suggests the importance of optimal levels of ZGRF1 for cell division. The extent of defects and proportion of microtubule-based defects observed were significantly higher in cells with the ZGRF1 variant proteins (Figure 3C, Supporting Figure S10). Mitotic defects in cells with ZGRF1 variant proteins during cell division and endogenous localization of the protein across mitotic stages suggest its role in cell division.

**Figure 3:**
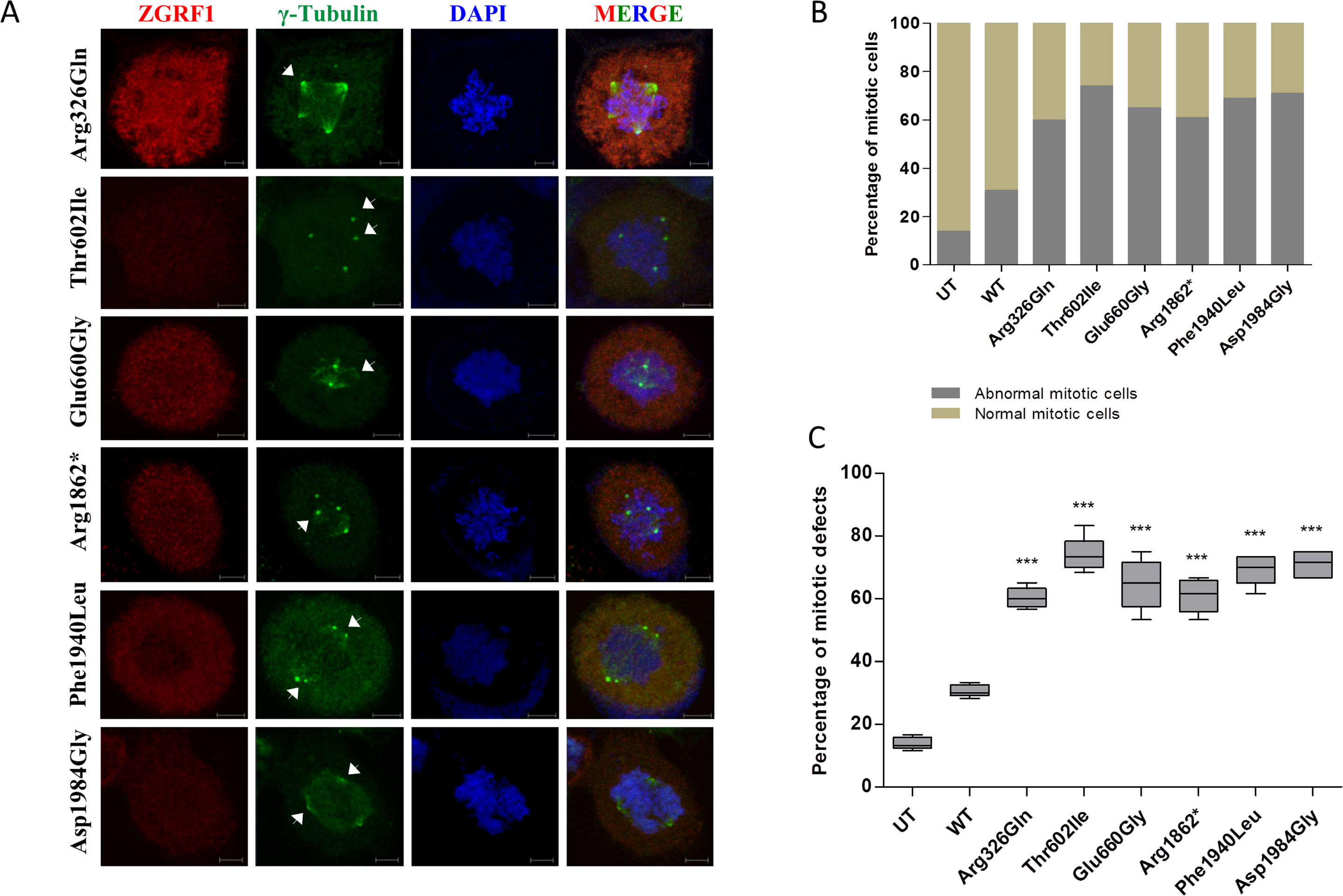
Mitotic defects in cells with ZGRF1 variants. (A) Defective cells with multipolar or disoriented spindles. Arrows point towards spindle poles. Scale bar = 5µm. (B) Percentage of normal and abnormal cells (with mitotic defects) in untransfected cells and cells with wild-type and six ZGRF1 variants (N=300). (C) Statistical analysis of the percentage of abnormal mitotic cells in untransfected HEK293 cells and cells with wild-type and six ZGRF1 variants (\*\*\**P* : ≤ 0.001).

### ZGRF1 and UPF1 co-localize

In search for potential homologs of ZGRF1 by *in-silico* analysis, we found that the C-terminal region of ZGRF1 shows similarities with the helicase/P-loop NTPase domain of DNA/RNA helicases bearing different types of zinc fingers. We obtained 13 candidates with high significance values, and from the highest similarity scores we found UPF1 (Up-frameshift 1) to be the nearest paralog of ZGRF1 (Figure 4A, Supporting Figure S11a). We then undertook pairwise alignment of ZGRF1 to the Protein Data Bank (PDB) UPF1-RNA complex (sequence ID: 2XZL, Chakrabarti et al., 2011), which shows 29% identity and 46% similarity between the two proteins (Supporting Figure S11b). UPF1 is a DNA/RNA helicase and a core protein of the nonsense-mediated mRNA decay (NMD) pathway. It is also involved in DNA repair, cell-cycle regulation and genome stability. ZGRF1 and UPF1 have similar protein domains at their C-terminal region. Both proteins contain the DNA2 and recD domains, and have zinc fingers and helicase regions comprised of P-loop NTPase domain. UPF1 has been reported to be a cytoplasmic protein (Applequist et al 1997). We observed that ZGRF1 and UPF1 are present in the cell cytoplasm, and localize together at the centrosomes, nucleolus, and spindle poles during mitosis (Figure 4B). In dividing cells, we found UPF1 to co-localize with γ-tubulin at the spindle poles and spindle fibres (Supporting Figure S11c).

**Figure 4:**
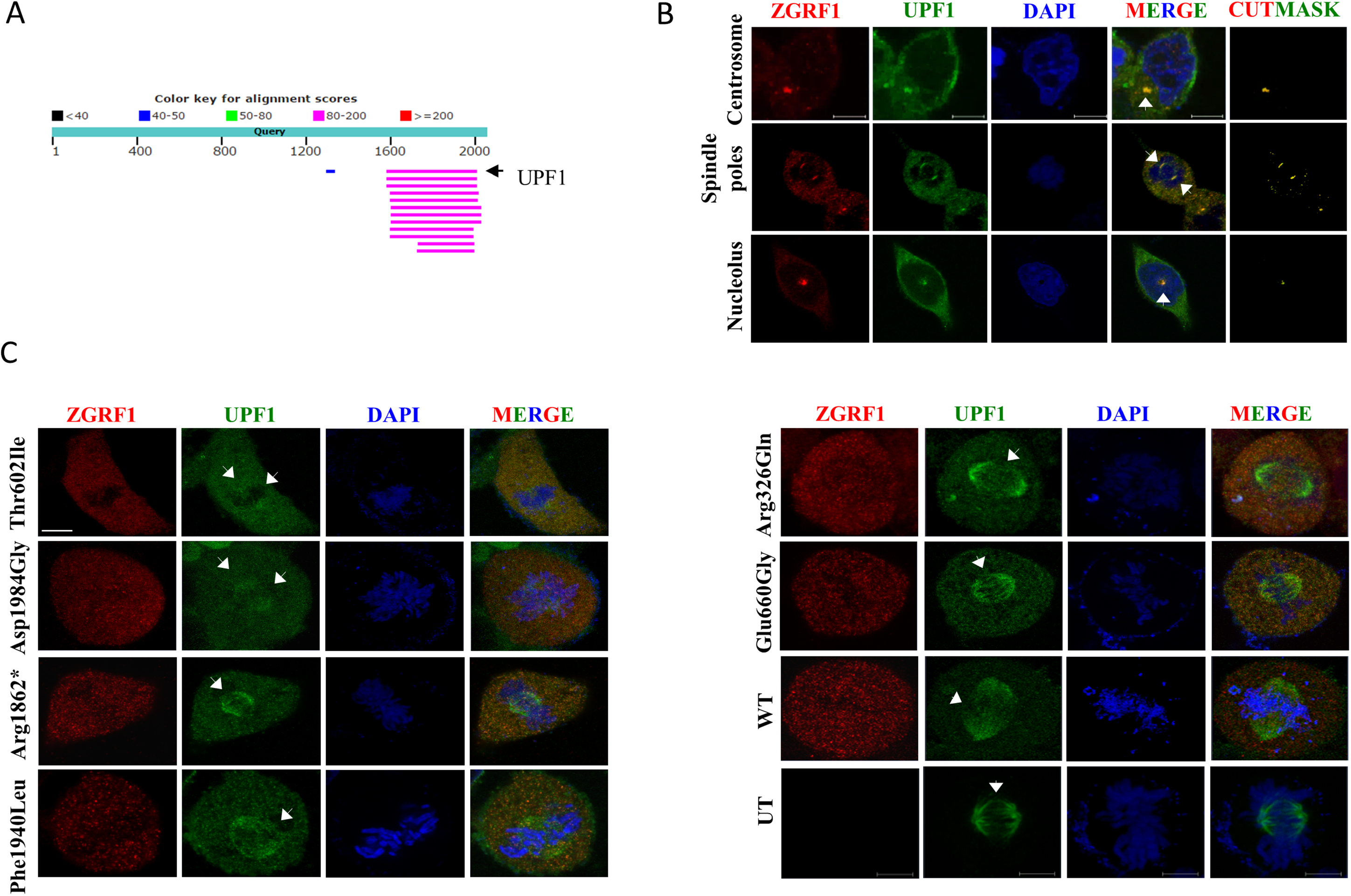
ZGRF1 and UPF1 expression. (A) BLAST alignment output for the ZGRF I full-length protein. (B) ZGRF1 and UPF1 in cultured HEK293 cells co-localize in the centrosomes, spindle poles and nucleolus indicated by arrows. Cutmask represents the regions of co-localization. Scale bar = 5 µm. (C) UPF1 localization to spindles was substantially disrupted by the Thr602Ile and Asp 1984Gly variants, partially disrupted by Argl862* and Phe1940Leu and unaffected by Arg326Glu and Glu660Gly. Arrows point towards spindle poles and microtubules. Scale bar = 5 µm.

### Partial disruption of UPF1 localization in cells with ZGRF1 variants

HEK293 cells transfected with the ZGRF1-wildtype and ZGRF1 variants were co-stained for ZGRF1 and UPF1. Considering that these two proteins co-localize in cellular compartments and at spindle poles during mitotic stages, it was of interest to examine impact, if any, on the UPF1 at spindles in the background of ZGRF1 wild-type or variant proteins. In the figure, only cells with two spindle poles have been shown to maintain consistency in data presentation (n=60). Cells with Thr602Ile and Asp1984Gly showed substantial disruption in UPF1 staining at the poles and microtubules (Figure 4C). For Arg1862* and Phe1940Leu, milder disruption was observed in the overall staining for UPF1 in the poles and spindles. However, Arg326Gln and Glu660Gly did not influence the localization pattern of UPF1. Taken together, these observations suggest that ZGRF1 associates with UPF1 and that Thr602Ile and Asp1984Gly negatively influence the localization of UPF1 in a dividing cell.

## Discussion

Though rare, cases of HWE have been reported from several parts of the world. Genetic causes underlying sensory reflex epilepsies are difficult to identify owing to complex inheritance, clinical-, genetic-, etiological-heterogeneity, and gene-environment interactions (Italiano et al., 2016). However, in recent years there has been progress in identifying eight genes/loci for light and sound induced epilepsies (Galizia et al., 2015; Kasteleijn-Nolst Trenité et al., 2015; Okudan and Özkara 2018).

While the cellular mechanisms underlying the manifestation of the disorder are yet to be understood, cortical networks in the brain in predisposed individuals have been indicated to play a role (Ferlazzo et al., 2005). Certain genetic mutations that alter the somatosensory networks could render individuals with these mutations susceptible to epileptic activity upon exposure to specific stimuli. Mutations in genes identified for HWE include: *SYN1*, involved in neurotransmitter release and synaptogenesis identified in a French Canadian family (Fassio et al., 2011; Nguyen et al., 2015); *GPR56*, a regulator of neural precursor cell migration identified in a Portuguese boy (Öncü-Öner et al., 2018; Santos-Silva et al., 2015) and *SLC1A1* which alters glutamate transport and predisposes to HWE, identified in Indian families (Karan et al., 2017).

We found six variants in ZGRF1 (p.Arg326Gln, p.Thr602Ile, p.Glu660Gly, p.Arg1862*, p.Phe1940Leu and p.Asp1984Gly) in HWE patients, including one (p.Thr602Ile) in the family (Family 227) where the locus was previously mapped (*HWE2*; Ratnapriya et al., 2009b). *HWE2* is one among the four known loci for HWE, the three others being 10q21-q22 (*HWE1*, MIM: 613339; Ratnapriya et al., 2009a), 9p23-p24 (Karan et al., 2018) and 8p12-p23 (Karan et al., unpublished). *ZGRF1* is a mostly unexamined gene, and therefore we gathered clues about its potential cellular roles from the limited amount of literature available and *in-silico* analysis. Potential mutations in ZGRF1 have been found in other neurological and psychiatric disorders: p.Leu48Met and p.Glu1363Lys were reported in a multi-generation family with childhood apraxia of speech (Peter et al., 2016), and p.Met502Val was identified in a large-scale exome study of schizophrenic individuals (Need et al., 2012). In the recently released large whole exome-based genetic study of epilepsy patients (Epi25 collaborative, 2019), *ZGRF1* (reported as *C4orf21*) was listed among the top 200 candidate genes with burden for ultra-rare variants enriched for damaging missense variants. Among the six rare variants identified in this study, the variant c.1805C>T (p.Thr602Ile) was observed among 2/18340 cases, one of GGE (generalized genetic epilepsy) and one of non-acquired focal epilepsy (NAFE), and absent among 16872 controls.

*In-silico* analysis of ZGRF1 has suggested three major domains in the protein: DUF2439, zf-GRF and helicase. DUF2439 has been implicated in meiotic chromosome segregation, crossover recombination and telomere maintenance in yeast (Polakova et al., 2016; Silva et al., 2016). There are at least six other zinc finger GRF-type proteins involved in aspects of DNA damage response, transcription or RNA regulatory activities. We found the helicase domain of ZGRF1 to be homologous to DNA/RNA helicases, with UPF1 (Up-frameshift 1) being its nearest paralog. UPF1 is a core protein of the nonsense-mediated mRNA decay (NMD) pathway which is seen at high levels in the hippocampus of patients with intractable temporal lobe epilepsy and in mice post status epilepticus (Mooney et al., 2017). We observed ZGRF1 and UPF1 to localize in the nucleolus, centrosomes and spindle poles in cultured mammalian cells. Interestingly, UPF1 localization at the spindles is substantially affected by p.Thr602Ile and p.Asp1984Gly, and partially by p.Arg1862* and p.Phe1940Leu. This suggests that the two proteins interact, and that ZGRF1 plays molecular roles similar to that of UPF1. ZGRF1 also belongs to the family of UPF1-like RNA helicases that are involved in aspects of DNA replication and repair, transcription regulation, RNA metabolism, genome integrity or topoisomerase activities. Mutations in genes belonging to this family have been implicated in neurological conditions like distal spinal muscular atrophy (*IGHMBP2*; Grohmann et al., 2001), spinocerebellar ataxia and amyotrophic lateral sclerosis (*SETX*; Fogel et al., 2014).

Cellular localization studies show ZGRF1 to be prominently present at the centrosomes. Further, it was seen to localize with γ-tubulin at the spindle poles, indicating a role for this protein in the cell cycle. ZGRF1 variants led to varying degrees of microtubule-based defects in the form of multipolar cells, disoriented spindles, misaligned chromosomes and abnormal cytokinesis during mitosis. The variants p.Thr602Ile, p.Phe1940Leu and p.Asp1984Gly caused significant (2.2-2.4 fold increase) mitotic defects compared to the wild-type protein. The other three variants p.Arg326Gln, p.Glu660Gly and p.Arg1862* caused a two-fold increase in such defects. It is of interest to note that centrosomal and cell cycle proteins like *CDKL5* (Barbiero et al., 2017) and *EFHC1* (de Nijs et al., 2012) have been previously reported for human epilepsy syndromes. *ZGRF1* variants identified in this study provide genetic evidence and preliminary insights into hitherto unanticipated roles of this protein associated with HWE. Functional consequences of the *ZGRF1* variants in cell-based assays indicate that subtle cellular defects in physiological conditions may contribute to the pathophysiology of this disorder.

Two previous reports identify *ZGRF1* as a candidate for DNA crosslink resistance (Smogorzewska et al., 2010) and regulator of mammalian homologous recombination (HR) machinery (Adamson et al., 2012). Our observations (data not shown) suggest cytoplasm-nucleus shuttling activity of ZGRF1 in response to DNA damage and its role in mRNA surveillance and decay. ZGRF1, a multi-domain and multi-functional protein, is an interesting addition to the diversity of already known proteins underlying sensory epilepsies, which include channel proteins, neurotransmitter regulators, receptors and DNA helicases. Further studies involving mice carrying null alleles and missense alleles found in HWE patients might provide insights into the roles of *ZGRF1* in the disorder.

## Web resources

The URLs for the web resources used in this study are:

RefSeq (https://www.ncbi.nlm.nih.gov/refseq/), Primer3 (http://bioinfo.ut.ee/primer3-0.4.0/), NCBI HomoloGene database (https://www.ncbi.nlm.nih.gov/homologene/), Clustal Omega (https://www.ebi.ac.uk/Tools/msa/clustalo//), SIFT (https://sift.bii.a-star.edu.sg/), PolyPhen-2 (http://genetics.bwh.harvard.edu/pph2/), MutationTaster (http://www.mutationtaster.org/), and FATHMM (http://fathmm.biocompute.org.uk/), DELTA-BLAST and pBLAST (https://blast.ncbi.nlm.nih.gov/Blast.cgi?PROGRAM=blastp&PAGE_TYPE=BlastSearch&LINK_LOC=blasthome), EMBOSS Needle (https://www.ebi.ac.uk/Tools/psa/emboss_needle/), 1000 Genomes (https://www.internationalgenome.org/1000-genomes-browsers/), Ensembl (https://asia.ensembl.org/index.html), Exome Sequencing Project (ESP) (https://evs.gs.washington.edu/EVS/), gnomAD (https://gnomad.broadinstitute.org/), NCBI Variation Viewer (https://www.ncbi.nlm.nih.gov/variation/view/).

## Acknowledgements

We thank the individuals with hot water epilepsy and their families for participation in this study. We thank Sambhavi Puri for her inputs and validation of the experiments comprising mitotic defects and Minu John for her help with sequencing experiments in the early stages. We thank Ram Shankar Mani, Nishtha Pandey, Praveen Raju, Kalpita Karan and Sharat Chandra for their comments on the manuscript. This work is supported by funds from ICMR, New Delhi; DBT, New Delhi and JNCASR, Bangalore. SRC acknowledges receipt of research fellowship from CSIR, New Delhi.

## Conflict of interest

All authors report that they have no financial interests or potential conflicts of interests.

## References

Adamson, B., Smogorzewska, A., Sigoillot, F. D., King, R. W., Elledge, S. J. (2012). A genome-wide homologous recombination screen identifies the RNA-binding protein RBMX as a component of the DNA-damage response. Nature Cell Biology, 14, 318–328. doi:10.1038/ncb2426

Adzhubei, I. A., Schmidt, S., Peshkin, L., Ramensky, V. E., Gerasimova, A., Bork, P., Kondrashov, A. S., Sunyaev, S. R. (2010). A method and server for predicting damaging missense mutations. Nature Methods, 7, 248–249. doi:10.1038/nmeth0410-248

Allen, I. M. (1945). Observations on cases of reflex epilepsy. The New Zealand Medical Journal, 44, 135–142

Applequist, S. E., Selg, M., Raman, C., Jäck, H. M. (1997). Cloning and characterization of HUPF1, a human homolog of the Saccharomyces cerevisiae nonsense mRNA-reducing UPF1 protein. Nucleic Acids Research, 25, 814–821. doi:10.1093/nar/25.4.814

Ashkenazy, H., Erez, E., Martz, E., Pupko, T., Ben-Tal, N. (2010). ConSurf 2010: calculating evolutionary conservation in sequence and structure of proteins and nucleic acids. Nucleic Acids Research, 38, W529–W533. doi:10.1093/nar/gkq399

Barbiero, I., Valente, D., Chandola, C., Magi, F., Bergo, A., Monteonofrio, L., Tramarin, M., Fazzari, M., Soddu, S., Landsberger, N., Rinaldo, C., Kilstrup-Nielsen, C. (2017). CDKL5 localizes at the centrosome and midbody and is required for faithful cell division. Scientific Reports, 7, 1–12. doi:10.1038/s41598-017-05875-z

Bebek, N., Gürses, C., Gokyigit, A., Baykan, B., Ozkara, C., Dervent, A. (2001). Hot water epilepsy: Clinical and electrophysiologic findings based on 21 cases. Epilepsia, 42, 1180–1184. doi:10.1046/j.1528-1157.2001.31000.x

Boratyn, G. M., Schäffer, A. A., Agarwala, R., Altschul, S. F., Lipman, D. J., Madden, T. L. (2012). Domain enhanced lookup time accelerated BLAST. Biology Direct, 7, 12. doi:10.1186/1745-6150-7-12

Chakrabarti, S., Jayachandran, U., Bonneau, F., Fiorini, F., Basquin, C., Domcke, S, Le Hir, H., Conti, E. (2011). Molecular Mechanisms for the RNA-Dependent ATPase Activity of Upf1 and Its Regulation by Upf2. Molecular Cell, 41, 693–703. doi:10.1016/j.molcel.2011.02.010

Cheng, Z., Muhlrad, D., Lim, M. K., Parker, R., Song, H. (2007). Structural and functional insights into the human Upf1 helicase core. The EMBO Journal, 26, 253–64. doi:10.1038/sj.emboj.7601464

de Nijs, L., Wolkoff, N., Coumans, B., Delgado-Escueta, A. V., Grisar, T., Lakaye, B. (2012). Mutations of EFHC1, linked to juvenile myoclonic epilepsy, disrupt radial and tangential migrations during brain development. Human Molecular Genetics, 21, 5106–5117. doi:10.1093/hmg/dds356

Engel J. (2001). A proposed diagnostic scheme for people with epileptic seizures and with epilepsy: report of the ILAE Task Force on Classification and Terminology. Epilepsia, 42, 796–803. doi:10.1046/j.1528-1157.2001.10401.x

Epi25 Collaborative. (2019). Ultra-Rare Genetic Variation in the Epilepsies: A Whole-Exome Sequencing Study of 17,606 Individuals. American Journal of Human Genetics, 05, 267–282. doi: 10.1016/j.ajhg.2019.05.020

Fassio, A., Patry, L., Congia, S., Onofri, F., Piton, A., Gauthier, J., Pozzi, D., Messa, M., Defranchi, E., Fadda, M., Corradi, A., Baldelli, P., Lapointe, L., St-Onge, J., Meloche, C., Mottron, L., Valtorta, F., Nguyen, D. K., Rouleau, G. A., Benfenati, F., Cossette, P. (2011). SYN1 loss-of-function mutations in autism and partial epilepsy cause impaired synaptic function. Human Molecular Genetics, 20, 2297–2307. doi:10.1093/hmg/ddr122

Ferlazzo, E., Zifkin, B. G., Andermann, E., Andermann, F. (2005). Cortical triggers in generalized reflex seizures and epilepsies. Brain, 128, 700–710. doi:10.1093/brain/awh446

Fogel, B. L., Cho, E., Wahnich, A., Gao, F., Becherel, O. J., Wang, X., Fike, F., Chen, L., Criscuolo, C., De Michele, G., Filla, A., Collins, A., Hahn, A. F., Gatti, R. A., Konopka, G., Perlman, S., Lavin, M. F., Geschwind, D. H., Coppola, G. (2014). Mutation of senataxin alters disease-specific transcriptional networks in patients with ataxia with oculomotor apraxia type 2. Human Molecular Genetics, 23, 4758–4769. doi:10.1093/hmg/ddu190

Frickey, T., & Lupas, A. N. (2004). Phylogenetic analysis of AAA proteins. Journal of Structural Biology, 146, 2–10. doi:10.1016/j.jsb.2003.11.020

Galizia, E. C., Myers, C. T., Leu, C., de Kovel, C. G. F., Afrikanova, T., Cordero-Maldonado, M. L., Martins, T. G., Jacmin, M., Drury, S., Chinthapalli, V. K., Muhle, H., Pendziwiat, M., Sander, T., Ruppert, A., Møller, R. S, Thiele, H., Krause, R., Schubert, J., Lehesjoki, A., et al. (2015). CHD2 variants are a risk .factor for photosensitivity in epilepsy. Brain, 138, 1198–1207. doi:10.1093/brain/awv052

Grohmann, K., Schuelke, M., Diers, A., Hoffmann, K., Lucke, B., Adams, C., Bertini, E., Leonhardt-Horti, H., Muntoni, F., Ouvrier, R., Pfeufer, A., Maldergem, L. V., Wilmshurst, J. M., Wienker, T. F., Sendtner, M., Rudnik-Schöneborn, S., Zerres, K., Hübner, C. (2001). Mutations in the gene encoding immunoglobulin μ-binding protein 2 cause spinal muscular atrophy with respiratory distress type 1. Nature Genetics, 29, 75–77. doi:10.1038/ng703

Italiano, D., Striano, P., Russo, E., Leo, A., Spina, E., Zara, F., Striano, S., Gambardella, A., Labate, A., Gasparini, S., Lamberti, M., De Sarro, G., Aguglia, U., Ferlazzo, E. (2016). Genetics of reflex seizures and epilepsies in humans and animals. Epilepsy Research, 121, 47–54. doi:10.1016/j.eplepsyres.2016.01.010

Iyer, L. M., Leipe, D. D., Koonin, E. V., Aravind, L. (2004). Evolutionary history and higher order classification of AAA+ ATPases. Journal of Structural Biology, 146, 11–31. doi:10.1016/j.jsb.2003.10.010

Karan, K. R., Satishchandra, P., Sinha, S., Anand, A.(2018). A genetic locus for sensory epilepsy precipitated by contact with hot water maps to chromosome 9p24.3-p23. Journal of Genetics, 97, 391–398.

Karan, K. R., Satishchandra, P., Sinha, S., Anand, A. (2017). Rare *SLC1A1* variants in hot water epilepsy. Human Genetics, 136, 693–703. doi:10.1007/s00439-017-1778-7

Kasteleijn-Nolst Trenité, D. G. A., Volkers, L., Strengman, E., Schippers, H. M., Perquin, W., de Haan, G., Gkountidi, A. O., van’t Slot, R., van de Graaf, S. F., Jocic-Jakubi, B., Capovilla, G., Covanis, A., Parisi, P., Veggiotti, P., Brinciotti, M., Incorpora, G., Piccioli, M., Cantonetti, L., Berkovic, S. F., Scheffer, I. E., et al (2015). Clinical and genetic analysis of a family with two rare reflex epilepsies. Seizure, 29, 90–96. doi:10.1016/j.seizure.2015.03.020

Keipert, J. A. (1969). Epilepsy precipitated by bathing: water-immersion epilepsy. Journal of Paediatrics and Child Health, 5, 244–247. doi:10.1111/j.1440-1754.1969.tb02759.x

Kumar, P., Henikoff, S., Ng, P. C. (2009). Predicting the effects of coding non-synonymous variants on protein function using the SIFT algorithm. Nature Protocols, 4, 1073–1081. doi:10.1038/nprot.2009.86

Kurata, S. (1979). Epilepsy precipitated by bathing: a follow-up study. Brain and Development (Domestic ed), 11, 400–405.

Li, H., & Durbin, R. (2009). Fast and accurate short read alignment with Burrows-Wheeler transform. Bioinformatics, 25, 1754–1760. doi:10.1093/bioinformatics/btp324

Li, H., Handsaker, B., Wysoker, A., Fennell, T., Ruan, J., Homer, N., Marth, G., Abecasis, G., Durbin, R., 1000 Genome Project Data Processing Subgroup. (2009). The Sequence Alignment/Map format and SAMtools. Bioinformatics, 25, 2078–2079. doi:10.1093/bioinformatics/btp352

Mani, K. S., Mani, A. J., Ramesh, C. K. (1974). Hot-water epilepsy-a peculiar type of reflex epilepsy: clinical and EEG features in 108 cases. Transactions of the American Neurological Association, 99, 224–226.

Mooney, C. M., Jimenez-Mateos, E. M., Engel, T., Mooney, C., Diviney, M., Venø, M. T., Kjems, J., Farrell, M. A., O’Brien, D. F., Delanty, N., Henshall, D. C. (2017). RNA sequencing of synaptic and cytoplasmic Upf1-bound transcripts supports contribution of nonsense-mediated decay to epileptogenesis. Scientific Reports, 7, 41517. doi:10.1038/srep41517

Moran, J. (1976). So-called water immersion epilepsy. Irish Journal of Medical Science, 145, 140

Morimoto, T., Hayakawa, T., Sugie, H., Awaya, Y., Fukuyama, Y. (1985). Epileptic Seizures Precipitated by Constant Light, Movement in Daily Life, and Hot Water Immersion. Epilepsia, 26, 237–242. doi:10.1111/j.1528-1157.1985.tb05412.x

Need, A. C., McEvoy, J. P., Gennarelli, M., Heinzen, E. L., Ge, D., Maia, J. M., Shianna, K. V., He, M., Cirulli, E. T., Gumbs, C. E., Zhao, Q., Campbell, C. R., Hong, L., Rosenquist, P., Putkonen, A., Hallikainen, T., Repo-Tiihonen, E., Tiihonen, J., Levy, D. L., Meltzer, H. Y., Goldstein, D. B. (2012). Exome Sequencing Followed by Large-Scale Genotyping Suggests a Limited Role for Moderately Rare Risk Factors of Strong Effect in Schizophrenia. American Journal of Human Genetics, 91, 303–312. doi:10.1016/j.ajhg.2012.06.018

Nguyen, D. K., Rouleau, I., Sénéchal, G., Ansaldo, A. I., Gravel, M., Benfenati, F., Cossette, P. (2015). X-linked focal epilepsy with reflex bathing seizures: Characterization of a distinct epileptic syndrome. Epilepsi., 56, 1098–1108. doi:10.1111/epi.13042

Okudan, Z. V. & Özkara, Ç. (2018). Reflex epilepsy: triggers and management strategies. Neuropsychiatric Disease and Treatment, 14, 327–337. doi:10.2147/NDT.S107669

Öncü-Öner, T., Ünalp, A., Porsuk-Doru, I., Agilkaya, S., Güleryüz, H.,, Saraç, A., Ergüner, B., Yüksel, B., Hız-Kurul, S., Cingöz, S. (2018). GPR56 homozygous nonsense mutation p.R271* associated with phenotypic variability in bilateral frontoparietalpolymicrogyria. The Turkish Journal of Pediatrics, 60, 229–237. doi:10.24953/turkjped.2018.03.001

Peter, B., Wijsman, E. M., Nato, A. Q., Matsushita, M. M., Chapman, K. L., Stanaway, I. B., Wolff, J., Oda, K., Gabo, V. B., Raskind, W. H. (2016). Genetic Candidate Variants in Two Multigenerational Families with Childhood Apraxia of Speech. PLOS One, 11, e0153864. doi:10.1371/journal.pone.0153864

Polakova, S., Molnarova, L., Hyppa, R. W., Benko, Z., Misova, I., Schleiffer, A., Smith, G. R., Gregan, J. (2016). Dbl2 Regulates Rad51 and DNA Joint Molecule Metabolism to Ensure Proper Meiotic Chromosome Segregation. PLOS Genetics, 12, e1006102. doi:10.1371/journal.pgen.1006102

Quinlan, A. R. & Hall, I. M. (2010). BEDTools: a flexible suite of utilities for comparing genomic features. Bioinformatics, 26, 841–842. doi:10.1093/bioinformatics/btq033

Raney, K. D., Byrd, A. K., Aarattuthodiyil, S. (2013). Structure and Mechanisms of SF1 DNA Helicases. Advances in Experimental Medicine and Biology, 767, 17–46. doi:10.1007/978-1-4614-5037-5_2

Ratnapriya, R., Satishchandra, P., Kumar, S. D., Gadre, G., Reddy, R., Anand, A. (2009a). A locus for autosomal dominant reflex epilepsy precipitated by hot water maps at chromosome 10q21.3-q22.3. Human Genetics, 125, 541–549. doi:10.1007/s00439-009-0648-3

Ratnapriya, R., Satishchandra, P., Dilip, S., Gadre, G., Anand, A. (2009b). Familial autosomal dominant refex epilepsy triggered by hot water maps to 4q24-q28. Human Genetics, 126, 677–683. doi:10.1007/s00439-009-0718-6

Santos-Silva, R., Passas, A., Rocha, C., Figueiredo, R., Mendes-Ribeiro, J., Fernandes, S., Biskup, S., Leão, M. (2015). Bilateral frontoparietalpolymicrogyria: A novel *GPR56* mutation and an unusual phenotype. Neuropediatrics, 46, 134–138. doi:10.1055/s-0034-1399754

Satishchandra, P. (2003). Geographically Specific Epilepsy Syndromes in India Hot-Water Epilepsy. Epilepsia, 44, 29–32. doi:10.1046/j.1528-1157.44.s.1.14.x

Satishchandra, P., Shivaramakrishana, A., Kaliaperumal, V. G., Schoenberg, B. S. (1988). Hot-Water Epilepsy: A Variant of Reflex Epilepsy in Southern India. Epilepsia, 29, 52–56. doi:10.1111/j.1528-1157.1988.tb05098.x

Schwarz, J. M., Rödelsperger, C., Schuelke, M., Seelow, D. (2010). MutationTaster evaluates disease-causing potential of sequence alterations. Nature Methods, 7, 575–576. doi:10.1038/nmeth0810-575

Shihab, H. A., Gough, J., Cooper, D. N., Stenson, P. D., Barker, G. L. A., Edwards, K. J., Day, I. N. M., Gaunt, T. R. (2013). Predicting the Functional, Molecular, and Phenotypic Consequences of Amino Acid Substitutions using Hidden Markov Models. Human Mutation, 34, 57–65. doi:10.1002/humu.22225

Silva, S., Altmannova, V., Luke-Glaser, S., Henriksen, P., Gallina, I., Yang, X., Choudhary, C., Luke, B., Krejci, L., Lisby, M. (2016). Mte1 interacts with Mph1 and promotes crossover recombination and telomere maintenance. Genes & Development, 30, 700–717. doi:10.1101/gad.276204.115

Smogorzewska, A., Desetty, R., Saito, T. T., Schlabach, M., Lach, F. P., Sowa, M. E., Clark, A. B., Kunkel, T. A., Harper, J. W., Colaiácovo, M. P., Elledge, S. J. (2010). A Genetic Screen Identifies FAN1, a Fanconi Anemia-Associated Nuclease Necessary for DNA Interstrand Crosslink Repair. Molecular Cell, 39, 36–47. doi:10.1016/j.molcel.2010.06.023

Stensman, R. & Ursing, B. (1971). Epilepsy precipitated by hot water immersion. Neurology, 21, 559–562. doi:10.1212/wnl.21.5.559

Szymonowicz, W. & Meloff, K. L. (1978). Hot Water Epilepsy. Canadian Journal of Neurological Sciences, 5, 247–251. doi:10.1017/s0317167100024616

