## Supplemental Data for "*ZGRF1* variants implicated in a sensory reflex epilepsy"

### Supporting Information

### Supporting Table S1: Primer sequences used to examine the *ZGRF1* transcript

| **Amplicon** | **Forward primer sequence5' → 3'** | **Amplicon** | **Reverse primer sequence5' → 3'** |
|---|---|---|---|
| ZGRF1-5'utr-a-F | tccggatcctgatagtctcg | ZGRF1-5'utr-a-R | tccttcccggttctcttctt |
| ZGRF1-5'utr-b+Ex1-F | ctccttgattcatgtctgtgg | ZGRF1-5'utr-b+Ex1-R | gctaaatctggcaacacagc |
| ZGRF1-Ex2-F | tccctccctctccttctctc | ZGRF1-Ex2-R | ccactgtgatatggggatca |
| ZGRF1-Ex3-F | tctcttaaacagactgatgc | ZGRF1-Ex3-R | acttaacctttctgcactgc |
| ZGRF1-Ex4-F | agtcttgcgattacgggtgt | ZGRF1-Ex4-R | aagtgccctttgaatgacaga |
| ZGRF1-Ex5i-F | ggcaaaccagaaacaatcctt | ZGRF1-Ex5i-R | ccttgtggttgtttttgagga |
| ZGRF1-Ex5ii-F | ctgaagtccgaatcatctag | ZGRF1-Ex5ii-R | catgtatttacctcctgagc |
| ZGRF1-Ex5iii-F | gcagtgcagaaaatgatggt | ZGRF1-Ex5iii-R | attcccatgtcaaaacccac |
| ZGRF1-Ex5iv-F | agggtgaacatttgccattc | ZGRF1-Ex5iv-R | gtgcttctgtgtctttggaa |
| ZGRF1-Ex5v-F | agtagtgacaacagtgtcca | ZGRF1-Ex5v-R | atggggacttttcattgcta |
| ZGRF1-Ex6-F | agtgctacattccagattga | ZGRF1-Ex6-R | agtcatgaaccaagatagct |
| ZGRF1-Ex7-F | gcccagccctagtatttctt | ZGRF1-Ex7-R | tgttagccaggatggtctct |
| ZGRF1-Ex8-F | ggcaagtgtgttttcacagg | ZGRF1-Ex8-R | gggggcaggagtcattaaa |
| ZGRF1-Ex9-F | ataggagaatttcccggttt | ZGRF1-Ex9-R | gagatattatgcttcttgtc |
| ZGRF1-Ex10-F | cccccaccatagctctacat | ZGRF1-Ex10-R | gcttcaagggatatgacactga |
| ZGRF1-Ex11a-F | gggacttcatcaaccctaagaa | ZGRF1-Ex11a-R | ttgcattgggatgtgttttg |
| ZGRF1-Ex11b-F | gcctgaggaccaaaatgaaa | ZGRF1-Ex11b-R | gccatctgtccaaattgctc |
| ZGRF1-Ex12-F | gggttaaaattgccattcg | ZGRF1-Ex12-R | gaagattgtaatttcccact |
| ZGRF1-Ex13-F | gactctgtttaaaggctattc | ZGRF1-Ex13-R | tcaagactcactctttaagc |
| ZGRF1-Ex14-F | ggagtacagtaatgccaagc | ZGRF1-Ex14-R | cagagcaagagcttgtaact |
| ZGRF1-Ex15-F | gatgacagttctgtgacagt | ZGRF1-Ex15-R | ggagataataaggctagactg |
| ZGRF1-Ex16-F | cagagcactcccaaccttta | ZGRF1-Ex16-R | ggaactcactgtacacaaca |
| ZGRF1-Ex17-F | gtttgggatctctgtgtgtg | ZGRF1-Ex17-R | ttacaggcatgagccactgc |
| ZGRF1-Ex18-F | cagttctcttctaccctcct | ZGRF1-Ex18-R | gggtagtcagttctgttctc |
| ZGRF1-Ex19-F | gtggaggttacagtgagctg | ZGRF1-Ex19-R | ccagcaagagtacatggaca |
| ZGRF1-Ex20-F | gccttcaagatggaataatgc | ZGRF1-Ex20-R | catggatgaagctggaaacc |
| ZGRF1-Ex21-F | tgacgagttaatgggtgcag | ZGRF1-Ex21-R | acatgcatgcaccttagtag |
| ZGRF1-Ex22-F | ttcttgttaatgggcttgtgg | ZGRF1-Ex22-R | aagctggtcgcaaactccta |
| ZGRF1-Ex23-F | tactcaggatacttcatggc | ZGRF1-Ex23-R | attatagggttccacttgtc |
| ZGRF1-Ex24+25-F | catggttgtaattcatctcc | ZGRF1-Ex24+25-R | tcaggcaccaaaccaactac |
| ZGRF1-Ex26-F | acagtggttgctttgagcag | ZGRF1-Ex26-R | agttacttcccccagcagag |
| ZGRF1-Ex27+3'utr-F | gctgggggaagtaactgaca | ZGRF1-Ex27+3'utr-R | gttccaacagatgagttctgg |

### Supporting Table S2: *ZGRF1* cDNA primers

| Amplicon | Forward primer sequence5’ → 3’ | Amplicon | Reverse primer sequence5’ → 3’ |
|---|---|---|---|
| ZGRF1-1F | ggaaagccaagaatttattg | ZGRF1-1R | tacatctttcttgccaacag |
| ZGRF1-2F | atatcctctggccgatctct | ZGRF1-2R | ctcagcacactcttcctgttg |
| ZGRF1-3F | gctgagatgaagagcacaga | ZGRF1-3R | tggagatgtttcagttctggc |
| ZGRF1-4F | gctcaggaggtaaatacatg | ZGRF1-4R | cagcatcagtccacttgaaag |
| ZGRF1-5F | tgtgggttttgacatgggaa | ZGRF1-5R | ctggcagtttcaacctcaaca |
| ZGRF1-6F | tccaaagacacagaagcaca | ZGRF1-6R | acctgcaagaagtcaatctgc |
| ZGRF1-7F | tcagaggacacagctcacag | ZGRF1-7R | aatgcatcccagaaacagcc |
| ZGRF1-8F | tccaggggatgtttcacttaa | ZGRF1-8R | gcaggttgactatgatggca |
| ZGRF1-9F | gcagactttcacatcttgcct | ZGRF1-9R | tccagctcaaagtctagggt |
| ZGRF1-10F | cagagaaaacagtatggcaa | ZGRF1-10R | aaatcaccactgccagcaag |
| ZGRF1-11F | tcagcctaggagcaacattga | ZGRF1-11R | tccaactactcgaacctgct |
| ZGRF1-12F | gaagacctgactcctacgga | ZGRF1-12R | tgaaagtccacagcactgag |
| ZGRF1-13F | gcaagtggaatagcaggctc | ZGRF1-13R | cttctgtttttcttccacttg |

### Supporting Table S3: Primer sets for site-directed mutagenesis

| **Primers** | **Primer sequence5' → 3'** |
|---|---|
| *ZGRF1*-Leu9X-sense | gaatttattgttctata**g**actcatcaaaagatgaagaa |
| *ZGRF1*-Leu9X-antisense | ttcttcatcttttgatgagt**c**tatagaacaataaattc |
| *ZGRF1*-Arg326Gln-sense | accatgagaaataaaagcc**a**gtgggccatgtatttatcc |
| *ZGRF1*-Arg326Gln-antisense | ggataaatacatggcccac**t**ggcttttatttctcatggt |
| *ZGRF1*-Thr602Ile-sense | gttagtgacaaacctacagtga**t**atttcctgttaaagagactctg |
| *ZGRF1*-Thr602Ile-antisense | cagagtctctttaacaggaaat**a**tcactgtaggtttgtcactaac |
| *ZGRF1*-Glu660Gly- sense | gatgctgtatacggagataataaag**g**agatgctaataaacctattcaa |
| *ZGRF1*-Glu660Gly-antisense | ttgaataggtttattagcatct**c**ctttattatctccgtatacagcatc |
| *ZGRF1*-Arg1862X- sense | gaaaatggattggaacaaactctttttgat**t**gactttgcttaatggg |
| *ZGRF1*-Arg1862X- antisense | cccattaagcaaagtc**a**atcaaaaagagtttgttccaatccattttc |
| *ZGRF1*-Phe1940Leu- sense | taatgtggcagaagctacg**c**ttacactcaagctgattc |
| *ZGRF1*-Phe1940Leu- antisense | gaatcagcttgagtgtaa**g**cgtagcttctgccacatta |
| *ZGRF1*-Asp1984Gly- sense | gtggactttcaccatcctg**g**tattaaaactgtgcaggtg |
| *ZGRF1*-Asp1984Gly- antisense | cacctgcacagttttaata**c**caggatggtgaaagtccac |

**Supporting Table S4: Summary statistics for the exome sequencing**

| Alignment features | Individual III:4 | | Individual IV:3 | |
|---|---|---|---|---|
| Read length in bases (paired) | 100 | | 100 | |
| Total number of reads[a] | 79978638 | | 84909810 | |
| Total reads for alignment (percentage high quality reads)[b] | 74872876 | (93.6) | 79634096 | (93.8) |
| Total reads aligned (percentage reads aligned)[c] | 74575432 | (99.60) | 79332908 | (99.62) |
| Reads aligned to whole exome (percentage reads aligned) | 51581554 | (69.17) | 54971128 | (69.29) |
| 4q24-q28 target length | 275107 | | 275107 | |
| 4q24-q28 target covered (percentage target covered) | 263491 | (95.78) | 263277 | (95.70) |

[a]Number of raw reads, [b] Number of reads yielded post processing of raw reads. Reads with >70% bases with Phred score >20, [c] All reads aligned, including exome and other regions of the genome.

**SupportingTable S5: Sequence coverage summary of the 4q24-q28 region**

| Coverage attributes | Individual III:4 | | Individual IV:3 | |
|---|---|---|---|---|
| | Chromosome 4q 24-q28 | Whole exome | Chromosome 4q24-q28 | Whole exome |
| %Total target covered with at least 5X read depth | 93.40 | 90.33 | 93.39 | 90.68 |
| %Total target covered with at least 10X read depth | 92.71 | 86.56 | 92.79 | 86.99 |
| %Total target covered with at least 15X read depth | 92.08 | 83.40 | 92.20 | 83.92 |
| %Total target covered with at least 20X read depth | 91.24 | 80.37 | 91.37 | 81.04 |

### Supporting Table S6: Twenty nine novel or rare variants with MAF ≤ 0.005 from the sequencing analysis of 4q24-q28

| Genomic position | Gene | Sequence variant | Hom in either or both (Y/-)[a] | Het and present in either sample (Y/-)[b] | Het and present in both samples (Y/-)[c] | Effect on protein | Variant ID | Frequency in control databases[d] | | | | Segregation (Y/N)[e] |
|---|---|---|---|---|---|---|---|---|---|---|---|---|
| | | | | | | | | ExAC | 1000G | ESP | dbSNP144 | |
| 103808458[g] | *CISD2* | NM_001008388.4:c.319-40T>C | - | Y | - | - | rs201682974 | 0.0010 | - | - | - | - |
| 103822082[g] | *SLC9B1* | NM_139173.3:c.*192C>T | - | - | Y | - | rs62327290 | - | - | - | - | N |
| 103822089[g] | *SLC9B1* | NM_139173.3:c.*185T>A | - | - | Y | - | rs62327291 | - | - | - | - | N |
| 103822298[g] | *SLC9B1* | NM_139173.3:c.1524G>A | - | - | Y | p.Gln508= | rs3175325 | 0.00003 | - | - | - | N |
| 103822492[g] | *SLC9B1* | NM_139173.3:c.1333-3C>T | - | - | Y | - | rs3974500 | 0.0029 | - | - | - | N |
| 103832539[g] | *SLC9B1* | NM_139173.3:c.936+49_936+50insC | - | - | Y | - | rs779644659 | 0.003 | - | - | - | N |
| 103832712[g] | *SLC9B1* | NM_139173.3:c.830-18delTA | - | - | Y | - | rs1553974206 | - | - | - | - | N |
| 104066176[g] | *CENPE* | NM_001813.2:c.4857+31_4857+32insA | - | - | Y | - | - | - | - | - | - | N |
| 104640282[f] | *TACR3* | NM_001059.2:c.548+3T>C | - | Y | - | - | - | - | - | - | - | - |
| 106155705[g] | *TET2* | NM_017628.4:c.606C>T | - | Y | - | p.Asn202= | rs201865755 | 0.0028 | 0.0038 | - | 0.0038 | - |
| 106068079[f] | *TET2* | NM_017628.4:c.-193+836_-193+839delAGAA | - | Y | - | - | rs572309712 | - | 0.0034 | - | 0.0034 | - |
| 106474096[f] | *ARHGEF38* | NM_017700:c.174C>T | Y | - | - | p.Thr58= | rs6533206 | 0.00007 | 0.0002 | - | 0.0002 | - |
| 107237722[g] | *AIMP1* | NM_001142416.1:c.18T>C | - | Y | - | p.Ala6= | - | - | - | - | - | - |
| 109543638[g] | *RPL34* | NM_033625.3:c.166-33C>T | - | Y | - | - | rs200789602 | 0.0008 | 0.0010 | 0.0002 | 0.0002 | - |
| 109578572[g] | *OSTC* | NM_021227.3:c.234-34G>A | Y | - | - | - | rs2851379 | 0.0044 | 0.0018 | 0.0047 | 0.0018 | - |
| 109578850[f] | *OSTC* | NM_021227.3:c.431+47T>C | - | Y | - | - | rs375863579 | 0.0023 | 0.0028 | 0.00007 | 0.0028 | - |
| 109767453[g] | *COL25A1* | NM_198721.3:c.1435-78_1435-77insT | Y | - | - | - | rs72620122 | - | - | - | - | - |

| Genomic position | Gene | Sequence variant | Hom in either or both (Y/-)[a] | Het and present in either sample (Y/-)[b] | Het and present in both samples (Y/-)[c] | Effect on protein | Variant ID | Frequency in control databases[d] | | | | Segregation (Y/N)[e] |
|---|---|---|---|---|---|---|---|---|---|---|---|---|
| | | | | | | | | ExAC | 1000G | ESP | dbSNP144 | |
| 110384047[g] | *SEC24B* | NM_006323.4:c.134-10_134-9insCTTTT | - | - | Y | - | rs70948092 | 0.0002 | - | 0.0164 | - | N |
| 110606395[f] | *CCDC109B* | NM_017918.4:c.817-13_817-12insT | -Y | Y | - | - | esv2223328 | - | - | - | - | - |
| 113539393[g] | *ZGRF1* | NM_018392.4:c.1805C>T | - | - | Y | p.Thr602Ile | rs201904239 | 0.0012 | 0.0012 | - | 0.0012 | Y |
| 113568126[g] | *LARP7* | NM_016648.3:c.552+15_552+16insA | - | - | Y | - | rs747753883 | - | - | - | - | N |
| 114267023[f] | *ANK2* | NM_020977.3:c.4249-33G>A | Y | - | - | - | rs9968405 | 0.0002 | 0.0002 | 0.0002 | 0.0002 | - |
| 114304735[f] | *ANK2* | NM_020977.3:c.*2108C>T | - | Y | - | - | rs190513908 | - | 0.0008 | - | 0.0008 | - |
| 114822867[f] | *ARSJ* | NM_024590.3:c.*561_*562insAA | - | Y | - | - | rs397732178 | - | - | - | - | - |
| 119035948[f] | *NDST3* | NM_0004784.2:c.1070-14_1070-13insC | Y | - | - | - | rs745946090 | 0.00004 | - | - | - | - |
| 123816124[f] | *FGF2* | NM_002006.4:c.*2573T>C | Y | - | - | - | rs2168915 | - | 0.0030 | - | 0.0030 | - |
| 123843750[f] | *NUDT6* | NM_145207.2:c.-548G>T | - | Y | - | - | rs200477292 | 0.0018 | 0.0024 | 0.0009 | 0.0024 | - |
| 123977466[g] | *SPATA5* | NM_145207.2:c.2080-76T>A | - | Y | - | - | rs558517923 | - | 0.0002 | - | 0.0002 | - |
| 124240031[f] | *SPATA5* | NM_145207.2:c.*4812G>C | - | Y | - | - | rs538982379 | - | 0.0026 | - | 0.0026 | - |

[a]Homozygous variants present in either or both of the exome sequenced individuals were not considered further.
[b]Heterozygous variants present in either of the two exome sequenced individuals were not considered further.
[c]Heterozygous variants present in both the exome sequenced individuals were confirmed by Sanger sequencing and analyzed for segregation in the family.
[d]Frequencies of all variants obtained from the analysis were determined in four databases. Variants with a MAF ≤ 0.005 in either of the databases were taken into consideration.
[e]Variants that were heterozygous in both the exome sequenced individuals and present in the proband, were examined for segregation in the family.
[f]Variants examined and eliminated by sequence analysis at >10X read depth in SAMtools.
[g]Variants confirmed and examined by Sanger sequencing.
Y = Yes, N = No, Hom = homozygous, Het = heterozygous

### Supporting Table S7: Primer sequences used to amplify variants with MAF≤0.005

| **Primers** | **Primer sequence5’ → 3’** |
|---|---|
| *CISD2*- c.319-40T>C-forward | tggttcagttggatgtagaatg |
| *CISD2*- c.319-40T>C-reverse | caaatacaaacagtagtgcagga |
| *SLC9B1*- c.*192C>T-forward | gggcctaaaatgcttacacg |
| *SLC9B1*- c.*192C>T-reverse | actggacatcatgggagttc |
| *SLC9B1*- c.*185T>A-forward | gggcctaaaatgcttacacg |
| *SLC9B1*- c.*185T>A-reverse | actggacatcatgggagttc |
| *SLC9B1*- c.1524G>A-forward | gggcctaaaatgcttacacg |
| *SLC9B1*- c.1524G>A-reverse | actggacatcatgggagttc |
| *SLC9B1*- c.1333-3C>T-forward | tttgctaggaagaacatggaac |
| *SLC9B1*- c.1333-3C>T- reverse | tcctggattctctgtacagtcc |
| *SLC9B1*- c.936+49_936+50insC-forward | tttgctaggaagaacatggaac |
| *SLC9B1*- c.936+49_936+50insC- reverse | tcctggattctctgtacagtcc |
| *SLC9B1*- c.830-18delTA-forward | ttcatgaaccacctagatttgc |
| *SLC9B1*- c.830-18delTA-reverse | ggcattggaggctagttttag |
| *CENPE*- c.4857+31_4857+32insA-forward | ttcaaggagcatcgcaaag |
| *CENPE*- c.4857+31_4857+32insA- reverse | gctgtcttctctgaaatattctgtc |
| *TET2*- c.606C>T-forward | tctgtgagttctgtagcccaag |
| *TET2*- c.606C>T-reverse | gtctgtgcggaattgatctg |
| *AIMP1*-c. 18T>C-forward | ggactgagcacaggaagagg |
| *AIMP1*-c. 18T>C-reverse | gcgaacggactacccattag |
| *RPL34*- c.166-33C>T-forward | ccaaatctgccacgaacatag |
| *RPL34*- c.166-33C>T-reverse | tcatttcctgctttgtgcag |
| *OSTC*- c.234-34G>A-forward | ggcacaaaaacggaaatagg |
| *OSTC*- c.234-34G>A-reverse | tcattgatgccaggacaaag |
| *COL25A1*-c. 1435-78_1435-77insT-forward | tgattgcatgtagaaggttgc |
| *COL25A1*-c. 1435-78_1435-77insT- reverse | catgccttacaatttgagttcc |
| *SEC24B*- c.134-10_134-9insCTTTT-forward | gggaggactacaagggtgtg |
| *SEC24B*- c.134-10_134-9insCTTTT-reverse | ggcagagtacatggctggac |
| *ZGRF1*- c.1805C>T-forward | agggtgaacatttgccattc |
| *ZGRF1*- c.1805C>T-reverse | gtgcttctgtgtctttggaa |
| *LARP7*- c.552+15_552+16insA-forward | tttgggaaatgtggcaatg |
| *LARP7*- c.552+15_552+16insA-reverse | cggcctttcttcttcttttc |
| *SPATA5*- c.2080-76T>A-forward | tgatgtgtggtttttggaatg |
| *SPATA5*- c.2080-76T>A-reverse | ttcggaaatgccaaagtctg |

### Supporting Figure S8

**a**

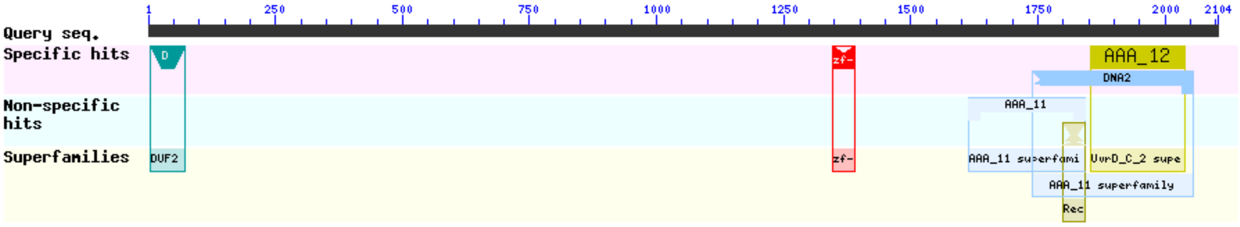


**b**

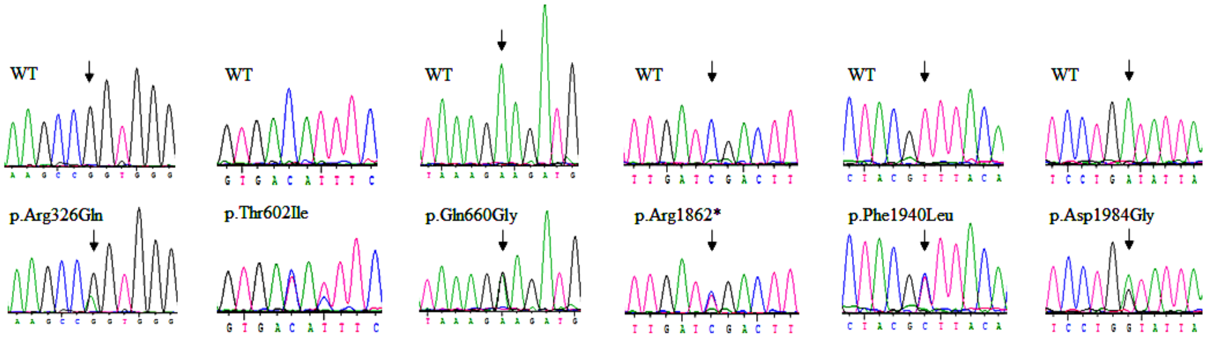


**Supporting Figure S8:** (a) Domains identified in by i*n-silico* analysis, (b) Representative electropherogram sequences of the *ZGRF1* variants in this study in the normal and affected individuals.

### Supporting Table S9: Polymorphisms found in *ZGRF1* in this study

| Nomenclature | Amino acid change | Ref SNP ID | Allele frequency in Indian controls (n=576) | Allele frequency in control databases |
|---|---|---|---|---|
| c.-284C>T | - | ≠ | 0.03 | - |
| c.-176C>A | - | ≠ | 0.01 | - |
| c.-53T>A | - | rs13104310 | 0.035 | 0.42 |
| c.22-99G>A | - | ≠ | 0.07 | - |
| c.102+26G>T | - | ≠ | 0.02 | - |
| c.142C>A | p.Leu48Met | rs61745597 | 0.012 | 0.0204 |
| c.162+13G>C | - | rs775648125 | 0.002 | 0.00012 |
| c.170C>T | p.Pro57Leu | rs775504533 | 0.02 | - |
| c.351+59G>T | - | rs1017468137 | 0.012 | - |
| c.351+28T>C | - | rs753126322 | 0.03 | 0.00034 |
| c.351+49G>T | - | ≠ | 0.01 | - |
| c.1035G>C | p.Gly345= | rs188866585 | 0.03 | 0.0005 |
| c.1071T>C | p.Asp357= | rs575464722 | 0.035 | 0.00012 |
| c.1229A>G | p.Asn410Ser | rs7696816 | 0.03 | 0.4137 |
| c.1677G>A | p.Glu557Lys | ≠ | 0.02 | - |
| c.1087T>C | p.Phe603Leu | rs78381215 | 0.02 | 0.026 |
| c.2802+92T>C | - | rs62316615 | 0 | 0.098 |
| c.3060A>G | p.Val1020= | ≠ | 0.01 | - |
| c.4434C>T | p.Ser1478= | rs561737889 | 0.005 | 0.0002 |
| c.4439-91G>A | - | rs186067473 | 0.001 | 0.0004 |
| c.4439-73A>G | - | ≠ | 0.022 | - |
| c.4439-41G>A | - | rs182036970 | 0.012 | 0.008 |
| c.4697+81G>T | - | rs6838559 | 0.022 | 0.4284 |
| c.5289T>C | p.Tyr1763= | rs141688719 | 0.0012 | 0.003 |
| c.5474+83C>G | - | rs78439826 | 0.007 | 0.022 |
| c.5475G>A | p.Arg1825= | rs78336146 | 0.026 | 0.0262 |
| c.5816C>T | p.Thr1939Met | rs150416544 | 0.012 | 0.0004 |
| c.*17T>G | - | ≠ | 0 | - |
| c.*52A>G | - | rs564397944 | 0.012 | 0.001 |

**Supporting Figure S10**

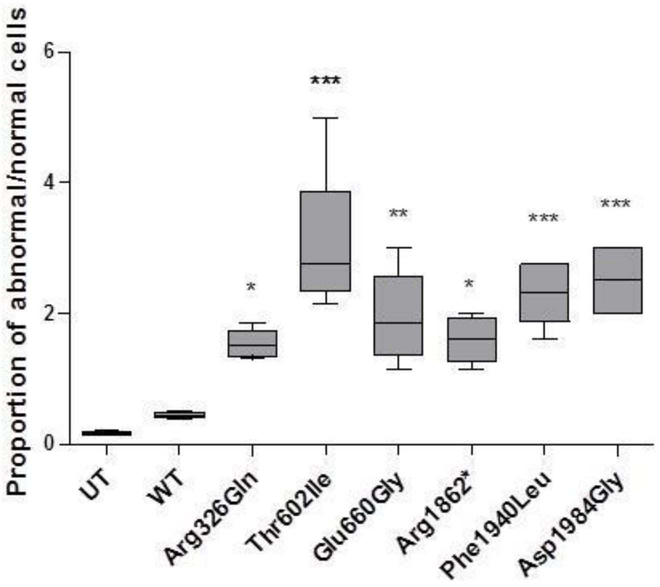


**Supporting Figure S10:** Proportion of abnormal and normal mitotic cells observed for the wild-type *ZGRF1*and variant ZGRF1 proteins (**P* ≤0.05, ***P* ≤0.01, and ****P* ≤0.001).

### Supporting Figure S11

**a**

**Sequences producing significant alignments:**

Select: All None Selected:0

Alignments Download GenPept Graphics Distance tree of results Multiple alignment

| Description | Max score | Total score | Query cover | E value | Ident | Accession |
|---|---|---|---|---|---|---|
| Chain A, Structural And Functional Insights Into The Human Upf1 Helicase Core | 197 | 197 | 20% | 1e-52 | 32% | 2GJK_A |
| Chain A, Upf1 Helicase - Rna Complex | 197 | 197 | 20% | 1e-52 | 32% | 2XZO_A |
| Chain A, Crystal Structure Of The Complex Between Human Nonsense Mediated Decay Factors Upf1 And Upf2 | 197 | 197 | 20% | 2e-51 | 32% | 2WJV_A |
| Chain A, Upf1-Rna Complex | 187 | 187 | 20% | 2e-48 | 33% | 2XZL_A |
| Chain A, Helicase Sen1 | 153 | 153 | 20% | 2e-37 | 29% | 5MZN_A |
| Chain A, Crystal Structure Of Dna2 Nuclease-helicase | 134 | 134 | 20% | 3e-31 | 26% | 5EAW_A |
| Chain A, Crystal Structure Of Dna2 In Complex With A 5' Overhang Dna | 134 | 134 | 20% | 3e-31 | 26% | 5EAN_A |
| Chain A, Crystal Structure Of Dna2 In Complex With An Ssdna | 134 | 134 | 20% | 4e-31 | 26% | 5EAX_A |
| Chain A, Crystal Structure Of Ighmbp2 Helicase In Complex With Rna | 126 | 126 | 19% | 4e-29 | 30% | 4B3G_A |
| Chain X, Crystal Structure Of 1ghmbp2 Helicase | 126 | 126 | 19% | 4e-29 | 30% | 4B3F_X |
| Chain A, Structural Insight Into The Function And Evolution Of The Spliceosomal Helicase Aquarius, Structure Of Aquarius In Complex With Amppnp | 87.4 | 87.4 | 13% | 1e-16 | 23% | 4PJ3_A |
| Chain U, Cryo-em Structure Of A Human Spliceosome Activated For Step 2 Of Splicing (c* Complex) | 87.0 | 87.0 | 13% | 2e-16 | 23% | 5MQF_U |
| Chain A, Crystal Structure Of Xenopus Laevis Apex2 C-terminal Znf-grf Domain | 47.0 | 47.0 | 2% | 5e-06 | 49% | 5U6Z_A |

**b**

Chain A, Upf1-Rna Complex

Sequence ID: 2XZL_A Length: 802 Number of Matches: 1

Range 1: 360 to 795 GenPept Graphics Next Match Previous Match

| Score | Expect | Method | Identities | Positives | Gaps |
|---|---|---|---|---|---|
| 415 bits(1067) | 4e-125 | Composition-based stats. | 138/471(29%) | 219/471(46%) | 55/471(11%) |

```
Query  1534  KLNKDQATALIQIAQMMASHESIEEVKELQTHTFPITIIHGVFGAGKSYLLAVVILFFVQ  1593
             +LN  Q+ A+  + Q                   P+++I G  G GK+   A ++
Sbjct  360   QLNSSQSNAVSHVLQR------------------PLSLIQGPPGTGKTVTSATIVYHL--  399

Query  1594  LFEKSEAPTIGNARPWKLLISSSTNVAVDRVLLGLLSLGFENFIRVGSVR--KIAKPILP  1651
                             ++L+ + +NVAVD +   L        +R+ +     +   +
Sbjct  400   ----------SKIHKDRILVCAPSNVAVDHLAAKL-RDLGLKVVRLTAKSREDVESSVSN  448

Query  1652  YSLHAGSENESEQLKELHALMKEDLTPTERVYVRKSIEQHKLGTNRTLLKQVRVVGVTCA  1711
              +LH      ++   +    +K+++         K   +    T   +L +  VV  TC
Sbjct  449   LALHNLVGRGAKGELKNLLKLKDEVG-ELSASDTKRFVKLVRKTEAEILNKADVVCCTCV  507

Query  1712  ACPFFPCMNDLKFPVVVLDECSQITEPASLLPIARFECEKLILVGDPKQLPPTIQGSDAAH  1771
                   ++  KF  V++DE +Q +EP  L+PI +   +++ILVGD +QL P I    AA
Sbjct  508   GAGDKRLDT-KFRTVLIDESTQASEPECLIPIVK-GAKQVILVGDHQQLGPVILERKAAD  565

Query  1772  ENGLEQTLFDRLCLMGHKPILLRTQYRCHPAISAIANDLFYKGALMNGVTEIERSPLLEW  1831
               GL+Q+LF+RL  +GH PI L  QYR +P +S   +++FY+G+L NGVT  +R+
Sbjct  566   -AGLKQSLFERLISLGHVPIRLEVQYRMNPYLSEFPSNMFYEGSLQNGVTIEQRTVPNSK  624

Query  1832  LP------TLCFYNVKGLEQ-IERDNSFHNVAEATFTLKLIQSLIASGIAGSMIGVITLY  1884
              P       + F+   G E+      SF N  EA    ++I  L   G+    IGVIT Y
Sbjct  625   FPWPIRGIPMMFWANYGREEISANGTSFLNRIEAMNCERIITKLFRDGVKPEQIGVITPY  684

Query  1885  KSQMYKLCHLLSAVDFHHPDIK-TVQVSTVDAFQGAEKEIIILSCVRTRQ---VGFIDSE  1940
             + Q   +   +        D+   V+V++VDAFQG EK+ IILSCVR  +   +GF+
Sbjct  685   EGQRAYILQYMQMNGSLDKDLYIKVEVASVDAFQGREKDYIILSCVRANEQQAIGFLRDP  744

Query  1941  KRMNVALTRGKRHLLIVGNLACLRKNQLWGRVIQH-------CEGREDGLQ  1984
             +R+NV LTR K  L+I+GN   L +N LW  ++ H        EG  D LQ
Sbjct  745   RRLNVGLTRAKYGLVILGNPRSLARNTLWNHLLIHFREKGCLVEGTLDNLQ  795
```

**c**

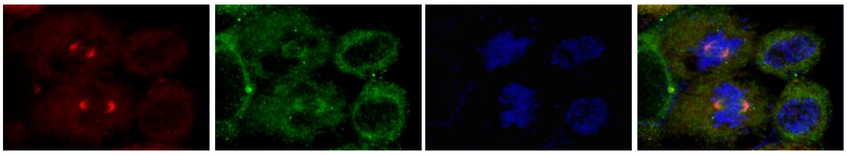


**Supporting Figure S11:** (a) Protein-BLAST search for ZGRF1 against Protein Data Bank (PDB) proteins found thirteen candidates with high similarity scores (b) Pairwise alignment of the ZGRF1 query sequence with the PDB entry 2XZL_A showing identity and similarity of 29% and 46%, respectively, between the amino acid residues (c) UPF1 co-localizes with γ-tubulin in dividing cells.
